# Redundant and specific roles of EGFR ligands in the ERK activation waves during collective cell migration of MDCK cells

**DOI:** 10.1101/2021.05.25.445569

**Authors:** Shuhao Lin, Daiki Hirayama, Gembu Maryu, Kimiya Matsuda, Naoya Hino, Eriko Deguchi, Kazuhiro Aoki, Ryo Iwamoto, Kenta Terai, Michiyuki Matsuda

## Abstract

Epidermal growth factor receptor (EGFR) plays a pivotal role in collective cell migration by mediating cell-to-cell propagation of extracellular signal-regulated kinase (ERK) activation. Here, we aimed to determine which EGFR ligands mediate the ERK activation waves by gene knockout. Four of the seven known EGFR ligands are expressed in MDCK cells. We found that epidermal growth factor (EGF)-deficient cells exhibited lower basal ERK activity than the cells deficient in heparin-binding EGF (HBEGF), transforming growth factor alpha (TGFα) or epiregulin (EREG), but all cell lines deficient in a single EGFR ligand retained the ERK activation waves. Therefore, we knocked out the EGFR ligand genes in decreasing order of expression. ERK activation waves were markedly suppressed, albeit incompletely, only when all four EGFR ligands were knocked-out. Re-expression of the EGFR ligands revealed that all but HBEGF could restore the ERK activation waves. Aiming at complete elimination of the ERK activation waves, we further attempted to knockout *Nrg1*, a ligand for ErbB3 and ErbB4, and found that *Nrg1* deficiency induced growth arrest in the absence of all four EGFR ligand genes expressed in MDCK cells. Collectively, these results showed that EGFR ligands exhibit remarkable redundancy in the propagation of ERK activation waves during collective cell migration.

## Introduction

Collective cell migration in mammalian tissues is a well-orchestrated cell movement underlying fundamental biological processes (Friedl and Gilmour, 2009; Mayor and Etienne-Manneville, 2016). The EGFR (epidermal growth factor receptor)-ERK (extracellular signal regulated kinase) signaling cascade plays a pivotal role in the collective cell migration of various cell types (Friedl and Gilmour, 2009; Yarden and Sliwkowski, 2001). During collective cell migration of epidermal cells, ERK activation propagates as multiple waves from the leader cells to the follower cells in an EGFR-dependent manner (Aoki et al., 2017; Boocock et al., 2021; Hino et al., 2020; Hiratsuka et al., 2015). Similar propagation of ERK activation waves has been visualized in developing Drosophila tracheal placode (Ogura et al., 2018), regenerating zebrafish scales (De Simone et al., 2021) and developing mouse cochlear ducts (Ishii et al., 2021), suggesting that growth factor-mediated ERK activation waves may generally underlie cell migration in various tissues.

EGFR is bound to and activated by a family of ligands that include epidermal growth factor (EGF), transforming growth factor alpha (TGFα), heparin-binding EGF-like growth factor (HBEGF), amphiregulin, betacellulin, epiregulin (EREG), and epigen (Harris et al., 2003). Extensive research has clarified the difference among the EGFR ligands with respect to binding affinity to four ErbB-family receptors including EGFR/ErbB1, sensitivity to proteases, subcellular localization, bioactivity to promote cell growth, and migration (Singh et al., 2016; Wilson et al., 2009). However, there still remain many questions to be answered about the roles played by the endogenous EGFR ligands, because much of the current knowledge is based on exogenous bolus application of EGFR ligands to tissue culture cells. Meanwhile, knockout mice deficient in each of the seven EGFR ligand genes are viable and fertile (Nanba et al., 2013; Schneider et al., 2008), suggesting functional redundancy among the EGFR ligands (Riese and Cullum, 2014; Singh and Coffey, 2014; Taylor et al., 2014; Zeng and Harris, 2014). Therefore, abrogation of multiple EGFR ligands is essential to clearly demonstrate the activity of the endogenous EGFR ligands.

Here, to determine which EGFR ligand mediates the propagation of the ERK activation waves during collective cell migration of MDCK cells, we knocked-out all four EGFR ligands expressed in MDCK cells, EGF, TGFα, HBEGF and EREG. We found that propagation of the ERK activation waves was markedly suppressed only when all four EGFR ligands were knocked-out. Re-expression of each EGFR ligand showed that EGF, TGFα and EREG, but not HBEGF, can restore the ERK activation waves.

## Results

### A new FRET biosensor for ERK without cross-reactivity to Cdk1

In previous studies (Aoki et al., 2017; Aoki et al., 2013; Hino et al., 2020), we used the FRET-based EKAREV biosensor for the detection of the ERK activation waves. However, EKAREV responds not only to ERK, but also to Cdk1, showing increases in FRET/CFP signal during mitosis even in the presence of an MEK inhibitor. To overcome this flaw, we developed a new ERK biosensor named EKARrEV by replacing the substrate peptide derived from Cdc25c with that from RSK1 (Fig. 1A). In MDCK cells expressing either EKAREV-NLS (with a nuclear localization signal) or EKARrEV-NLS, the FRET/CFP ratio was increased by EGF and decreased by the MEK inhibitor trametinib (Fig. 1B and C). In EKAREV-NLS-expressing cells, but not in EKARrEV-NLS-expressing cells, the FRET/CFP ratio increased immediately before mitosis (Fig. 1D). Although EKARrEV-NLS exhibited a smaller dynamic range and higher basal FRET/CFP ratio than the prototype EKAREV-NLS, we adopted EKARrEV-NLS because the increase in the FRET/CFP ratio during mitosis hampers automatic image analysis of ERK activation waves. It should be noted that Ponsioen et al. also recently reported another EKAREV-derived biosensor by modifying the substrate peptide to eliminate the cross-reactivity to Cdk1 (Ponsioen et al., 2021).

**Figure 1.**
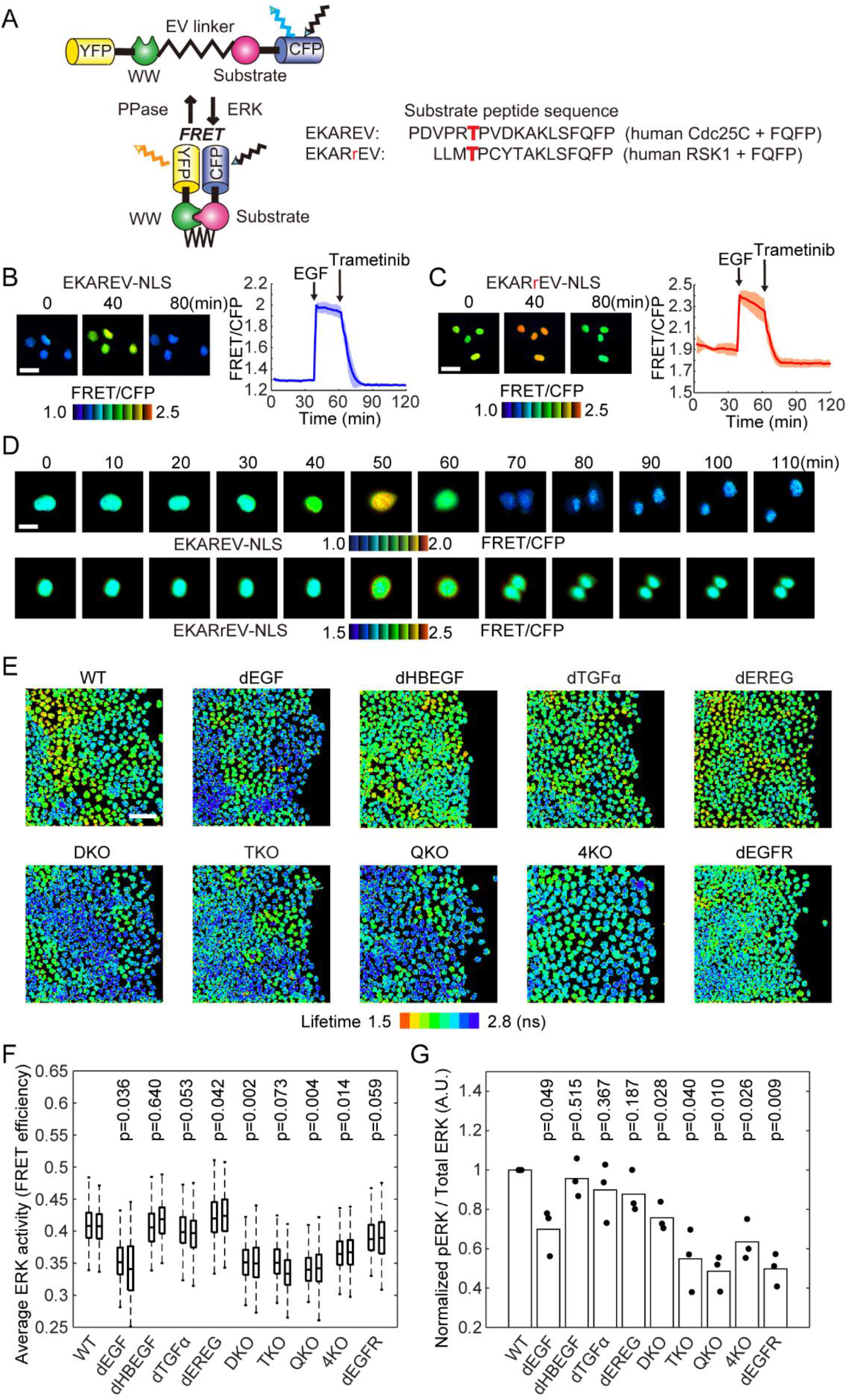
Properties of EKARrEV-NLS biosensor and ERK activity of MDCK cell lines after knockout of EGFR ligand genes. (A) Mode of action of the intramolecular FRET biosensors for ERK, EKAREV and EKARrEV. The red character T indicates the phosphorylation site. (B and C) MDCK cells expressing EKAREV-NLS (B) or EKARrEV-NLS (C) were observed under a wide field fluorescence microscope to acquired FRET/CFP time lapse images. During imaging, 10 ng mL^-1^ EGF and 200 nM trametinib were added. Representative ratio images are shown in the IMD mode. Line plots show time courses of the FRET/CFP ratio for 4 cells from a single experiment. Solid lines represent the means; shaded areas represent SD. Scale bar, 40 μm. (D) MDCK cells expressing EKAREV-NLS (top) or EKARrEV NLS (bottom) were imaged during mitosis, shown in the IMD mode. Scale bar, 20 μm. (E) Lifetime images of the donor fluorescence was acquired by a confocal microscope equipped with a 440 nm picosecond pulsed diode laser. MDCK cells are WT, single gene knockout (dEGF, dHBEGF, dTGFα, and dEREG), *Egf*/*Hbegf* double knockout (DKO), *Egf*/*Hbegf*/*Tgfα* triple knockout (TKO), and *Egf*/*Hbegf*/*Tgfα*/*Ereg* quadruple knockout (QKO and 4KO), and *Efgr* knockout (dEGFR). (F) Cells are timelapse imaged every 10 minutes for 100 min. FRET efficiency of randomly chosen cells was plotted for each cell line. The total number of analyzed cells from two independent experiments is as follows: WT, 271 and 231 cells; dEGF, 250 and 255 cells; dHBEGF, 260 and 136 cells; dTGFα, 227 and 163 cells; dEREG, 136 and 155 cells; DKO, 311 and 241 cells; TKO, 253 and 229 cells; QKO, 263 and 232 cells; 4KO, 252 and 255 cells; dEGFR, 217 and 155 cells. (G) Nine hours after the removal of silicone confinement, MDCK cells were analyzed by immunoblotting with anti-phospho-ERK or anti-pan-ERK antibody. The phospho-ERK signal normalized to the pan-ERK signal is shown. Data from three independent experiments are shown. p values of two-tailed t-test are shown on the top of panels F and G.

### Dependency on EGF for the mean ERK activity in serum-starved MDCK cells

Among the 7 known EGFR ligands, EGF, HBEGF, TGFα, and EREG were detected by RNA-Seq analysis of MDCK cells (Fig. S1) (Shukla et al., 2015). The expression of amphiregulin is marginal; therefore, we did not further pursue its role. To examine the roles of these EGFR ligands in the propagation of ERK activation waves, we employed CRISPR/Cas9-mediated knockout of each EGFR ligand gene (Fig. S2A). The resulting cell lines with knockout of a single EGFR ligand were named MDCK-dEGF, MDCK-dHBEGF, MDCK-dTGFα, and MDCK-dEREG, or simply dEGF, dHBEGF, dTGFα, and dEREG hereinafter. Because none of them exhibited a detectable decrease in the ERK activation waves, we knocked out the EGFR ligands sequentially from two abundantly expressed genes, *Egf* and *Hbegf*. Because the ERK activation waves were clearly visible in the double knockout cells, which we named DKO, *Tgfα* was further knocked out to obtain triple knockout cells, TKO, and then *Ereg* was knocked out to obtain quadruple knockout cells, QKO. After finding marked suppression of the ERK activation waves in QKO, all four genes were knocked out simultaneously to obtain an additional clone deficient from the four EGFR ligand genes, designated as 4KO. *Egfr* was also knocked-out for comparison, generating the dEGFR cell line (Fig. S2B). After single cell cloning, frame-shift mutations of both alleles were confirmed by genome sequencing (Fig. S2C).

We first examined the ERK activity using population-based methods. The mean FRET efficiency of the biosensor was determined for each cell line by fluorescence lifetime microscopy (Fig. 1E and F). Cells were also subjected to immunoblotting with anti-phospho-ERK antibody (Fig. 1G). In both experiments, cell lines deficient from *Egf*, dEGF, DKO, TKO, QKO, and 4KO exhibited lower ERK activity than the wild type (WT) cells. The level of ERK activity in these *Egf*-deficient cell lines was comparable to that in dEGFR cells, indicating that EGF is the principal endogenous EGFR ligand maintaining the basal EGFR activity in serum-starved MDCK cells. Of note, EKARrEV reflects the balance between ERK activity and phosphatase activity, whereas the anti-phospho-ERK antibody detects the phosphorylation of the catalytic loop of ERK. The discrepancy in the ERK activity of dEGFR cells between fluorescence lifetime imaging microscopy and immunoblotting may arise from this difference.

### Redundant roles of EGFR ligands in the propagation of ERK activation waves during collective cell migration as revealed by single cell analysis

For the analysis of collective cell migration, MDCK cells expressing EKARrEV-NLS were seeded one day before imaging within a culture-insert. After the removal of the culture-insert, cells were imaged in serum-free medium for more than 12 hours (Fig. 2A and Video 1). In this study, most experiments, unless noted otherwise, were performed in the absence of serum to exclude the effect of serum-derived growth factors. To evaluate the migration speed, cells that were located less than 100 µm from the leading edge at the time of confinement release were classified as the leader and submarginal cells (Fig. 2B). We analyzed these cells together because the leader cells are often replaced by the following submarginal cells during long-term imaging. After single cell tracking analysis, the displacement of the leader and submarginal cells before and 12 h after the start of collective cell migration was defined as the cell migration distance. The impairment of migration was clear when the two most abundant EGFR ligands, EGF and HBEGF, were absent, suggesting that the total amount of EGFR ligands may be the primary determinant for the migration of the leader and submarginal cells.

**Figure 2.**
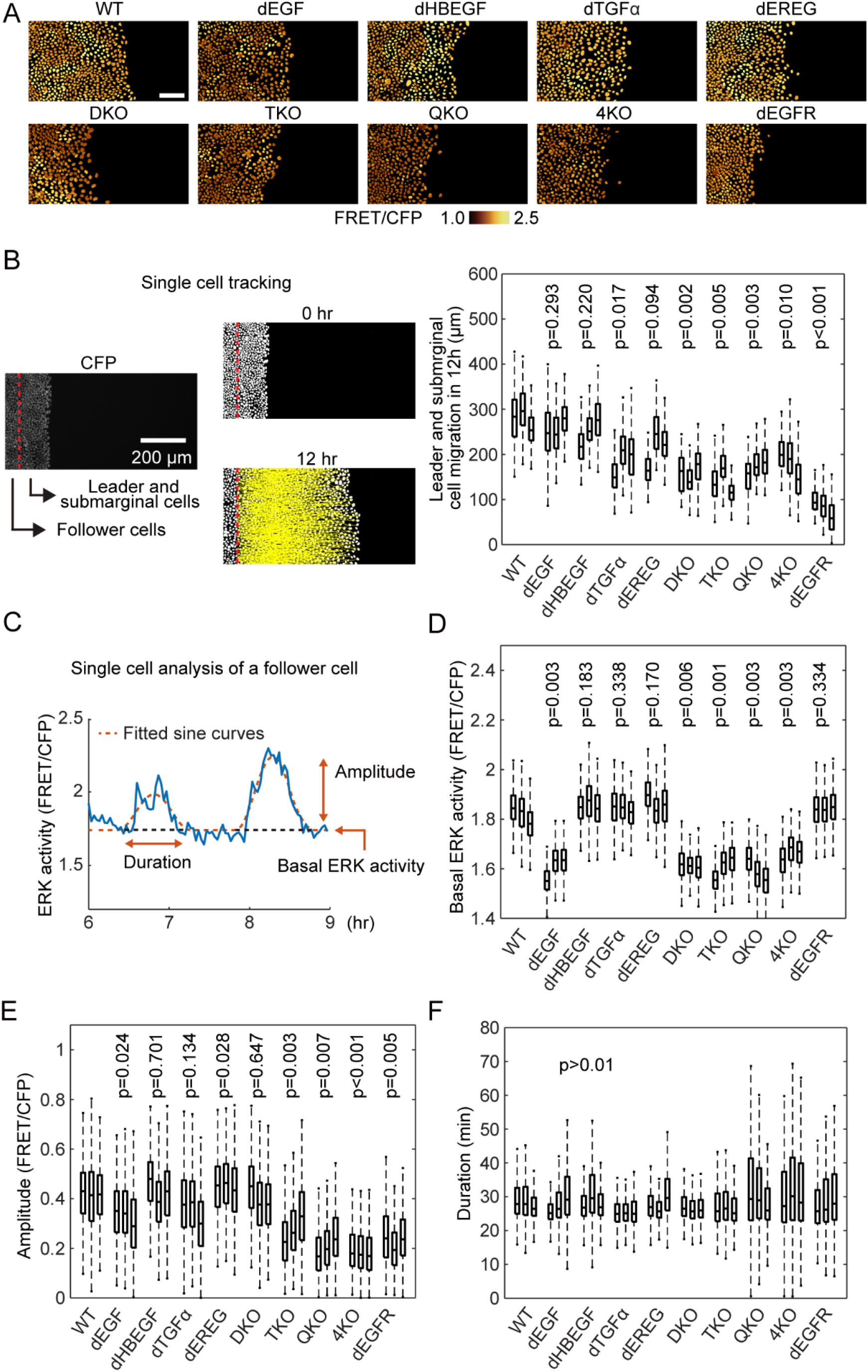
Single-cell analyses of ERK-activation dynamics in MDCK cells deficient of EGFR ligands. MDCK cells expressing EKARrEV were subjected to confinement release assay. (A) Time lapse images of FRET/CFP ratio were acquired every 2 minutes for up to twelve hours to generate Video 1. Shown are snap shots cropped from the Video. Scale bar, 100 μm. (B) We defined the cells locating within 100 µm from the edge of the open space as the leader and submarginal cells, the others as follower cells. CFP channel images were binarized for single cell detection. From the track of the leader and submarginal cells, migration distance in twelve hours was measured (left). Boxplot of leader and submarginal cell migration in twelve hours observation (right). The total number of analyzed cells from three independent experiments is as follows: WT, 395, 167, and 298 cells; dEGF, 386, 408 and 178 cells; dHBEGF, 447, 370 and 478 cells; dTGFα, 408, 366, and 388 cells; dEREG, 505, 525, and 504 cells; DKO, 256, 233, and 304 cells; TKO, 197, 270, and 229 cells; QKO, 278, 463 and 441 cells; 4KO, 407, 233 and 154 cells; dEGFR, 584, 303 and 217 cells. (C) After applying a mask for nuclei of the follower cells, the time course of FRET/CFP ratio in each cell was analyzed from six to nine hours after the start of migration (blue line). The ratio values were fitted by sine curves for the detection of waves (red line). For each wave, basal ERK activity (D), amplitude (E) and duration (F) were determined. The numbers of analyzed cells are as follows: WT, 814, 801, and 990 cells; dEGF, 789, 596 and 789 cells; dHBEGF, 846, 617 and 683 cells; dTGFα, 808, 762, and 838 cells; dEREG, 762, 1030, and 953 cells; DKO, 638, 656, and 560 cells; TKO, 649, 778, and 567 cells; QKO, 507, 638 and 567 cells; 4KO, 681, 291 and 451 cells; dEGFR, 643, 493 and 684 cells. p values of two-tailed t-test (WT to others) are shown on the top of panels B, D-F.

In WT cells, clear ERK activation waves were propagated from the leader cells to the follower cells (Video 1). Similarly, clear ERK activation waves were observed in single knockout cells, DKO, and even TKO cells. Only when all four EGFR ligand genes were knocked out in QKO and 4KO were the ERK activation waves markedly suppressed. To quantitatively understand the contribution of each EGFR ligand to the propagation of the ERK activation waves, we employed single cell-based analysis of the follower cells (Fig. 2C). After sine curve fitting of the time course of the FRET/CFP ratio in each cell (Hiratsuka et al., 2015), three parameters—basal ERK activity, amplitude, and duration—were obtained to evaluate the effect of EGFR ligand deficiency. Basal ERK activity was decreased in the *Egf-*deficient cell lines, dEGF, DKO, TKO, QKO and 4KO (Fig. 2D), as observed in the population-based analysis (Fig. 1E–G). The decrease of amplitude was obvious when all four EGFR ligand genes were knocked out (Fig. 2E). Negligible changes of duration period were observed in all cell lines, although the variance was markedly larger in QKO, 4KO, and dEGFR, probably because of the low amplitude of each pulse (Fig. 2F). These results indicated that EGF is the primary EGFR ligand to maintain the basal ERK activity, whereas all EGFR ligands may contribute to the propagation of ERK activation.

### Redundant roles of EGFR ligands in the propagation of ERK activation waves as revealed by heat map analysis

We then employed population-based analysis of ERK activation waves. First, a heat map of the ERK activity was obtained by interpolation of the FRET/CFP images (Fig. 3A). The directedness and the number of waves were analyzed by particle image velocimetry (PIV) and kymograph, respectively. We sometimes observed waves propagating in random directions, particularly at the area remote from the leader cells. PIV analysis showed that the directionality of ERK activation waves from the leader cells to the follower cells was markedly perturbed in QKO, 4KO and dEGFR cells (Fig. 3B). Similarly, the number of ERK activation waves was markedly decreased in QKO, 4KO, and dEGFR cells (Fig. 3C). Among the cell lines, we did not observe apparent changes in the speed of the ERK activation waves when they appeared.

**Figure 3.**
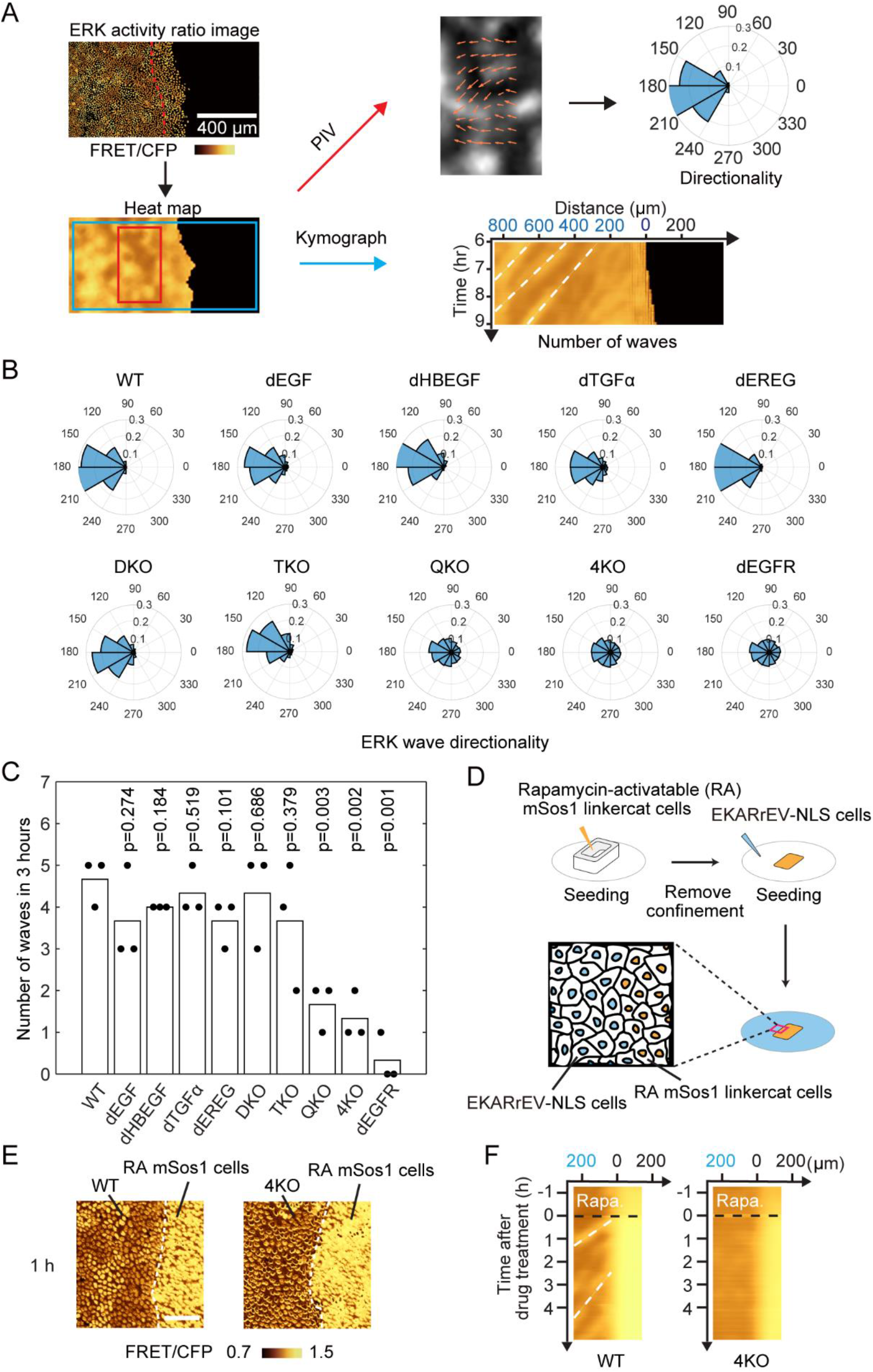
Propagation of ERK activation waves from leader cells. (A) Heat maps of ERK activity were obtained by interpolating the signals between the nuclei of cells and used for particle image velocimetry (PIV) and kymograph analysis. Directionality was measured by PIV and shown with polar histogram. Kymograph was obtained by interpolated ratio image. White broken lines indicate representative ERK waves. (B) The direction of ERK activation wave from six to nine hours after releasing of silicone confinement was shown by Polar histograms. Shown are summation of three independent experiments. (C) During the period of six to nine hours after the release of silicone confinement, the number of ERK activation waves from the leader cells was manually counted on the kymograph. Each dot represents the number of counted ERK activation waves in an independent experiment. p values of two-tailed t-test (WT to thers) are shown on the top. (D-F) For the artificial generation of ERK activation waves, MDCK cells expressing EKARrEV and rapamycin-activatable mSos1 was seeded in a silicone confinement. MDCK cells expressing EKARrEV alone were seeded in the surrounding area (D). Upon rapamycin addition, ERK activation waves were generated at the border of cell lines. (E) Time lapse FRET/CFP ratio images were acquired to generate Video 2. Shown are representative FRET/CFP ratio images 1 hour after addition of rapamycin. White broken lines indicate the border of cell lines. Scale bar, 50 μm. (F) Kymographs of ERK activity in MDCK-WT and MDCK-4KO cells upon rapamycin addition. White broken lines indicate representative ERK waves.

We previously showed that the ERK activation waves are induced by mechanical stretch from the leader cells, which retain high ERK activity (Hino et al., 2020). Therefore, the lack of ERK activation waves in QKO and 4KO cells may be due to the low ERK activity in the leader cells. To eliminate this possibility, we employed a rapamycin-activatable (RA) system combined with mSos1, a Ras guanine nucleotide exchange factor (Aoki et al., 2017). MDCK cells carrying RA-mSos1 were seeded next to MDCK WT or 4KO cells (Fig. 3D). Upon rapamycin treatment, ERK was activated in the RA-SOS cells, followed by the emergence of ERK activation waves in WT cells, but not in 4KO cells, at the interface (Fig. 3E, 3F and Video 2), indicating that ERK activation in RA-mSOS1 cells cannot be transmitted to the neighboring 4KO cells. In short, none of the knockouts of single EGFR ligands affected either the collective cell migration or the propagation of ERK activation waves, which were markedly impaired in cell lines deficient in all four EGFR ligand genes.

### Restoration of ERK activation waves by re-expression of the EGFR ligands

Do all EGFR ligands contribute to the ERK activation waves? To answer this question, we re-expressed each of the four EGFR ligands in 4KO cells and performed a confinement release assay (Fig. 4A and Video 3). The results showed that all EGFR ligands accelerated the migration of the leader and submarginal cells (Fig. 4B), but none of them increased the basal ERK activity to the level in the WT cells (Fig. 4C). Interestingly, except HBEGF, all EGFR ligands restored the ERK activation waves, as evidenced by the increase in the number, amplitude and directionality of waves from the leader cells (Fig. 4D–F). We also quantified the area of ERK activation waves by using binarized interpolated FRET/CFP ratio images (Fig. 4G). Again, except HBEGF, all EGFR ligands were able to restore the area of ERK activation waves.

**Figure 4.**
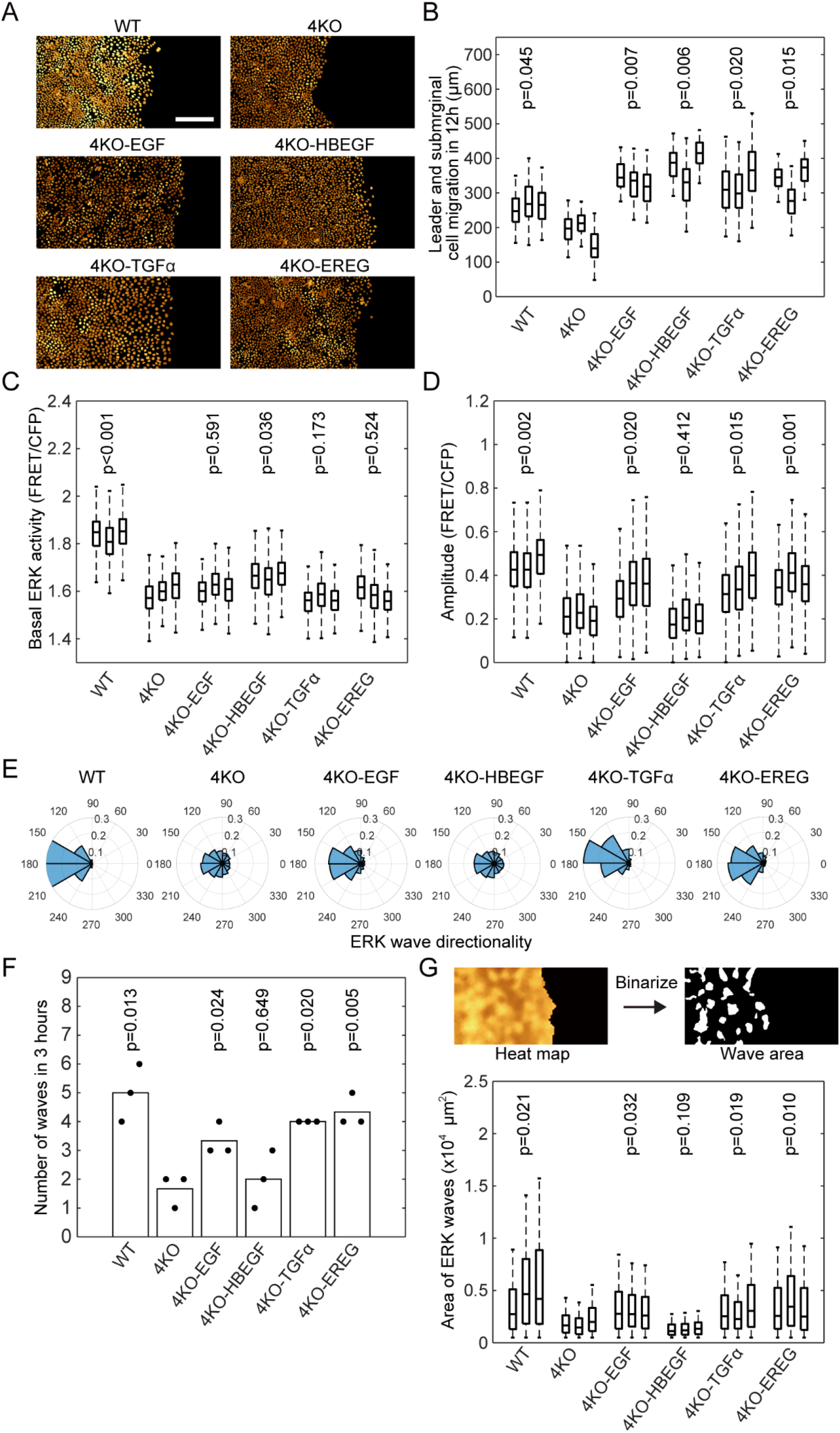
Restoration of ERK activation waves by expression of EGFR ligands in 4KO cells. The MDCK-4KO cells expressing the EKARrEV biosensor were transfected stably with an expression vector for cDNA of canine Egf, Hbegf, Tgfα, or Ereg; the resulting cell lines were named as 4KO-EGF, 4KO-HBEGF, 4KO-TGFα and 4KO-EREG, respectively. WT, 4KO and EGFR ligands expressing 4KO cells were subjected to confinement release assay. Three independent experiments were performed for each cell line. (A) Time lapse FRET/CFP ratio images were acquired to generate Video 3. Shown are snap shots cropped from the Video. Scale bar, 200 μm. (B) Boxplot of leader and submarginal cell migration in twelve hours observation. The number of analyzed cells from three independent experiments is as follows: WT, 357, 215 and 283 cells; 4KO, 348, 274, and 328 cells; 4KO-EGF, 130, 426, and 420 cells; 4KO-HBEGF, 224, 360, and 368 cells; 4KO-TGFα, 192, 300, and 215 cells; 4KO-EREG, 112, 404, and 373 cells. Data of three independent experiments are shown. (C-D) Box plots of basal ERK activity (C) and amplitude (D) of ERK activation in each cell line from six to nine hours after releasing of silicone confinement. The numbers of analyzed cells are as follows: WT, 703, 945 and 653 cells; 4KO, 656, 446, and 556 cells; 4KO-EGF, 576, 697, and 732 cells; 4KO-HBEGF, 667, 596, and 556 cells; 4KO-TGFα, 452, 638, and 484 cells; 4KO-EREG, 757, 786, and 902 cells. Data of three independent experiments are shown. (E) Polar histograms showing the distribution of ERK wave direction from six to nine hours after releasing of silicone confinement. Shown are summation of three independent experiments. (F) During the period of six to nine hours after the release of silicone confinement, the number of ERK activation waves from the leader cells was counted. Each dot represents the number of counted ERK activation waves in an independent experiment. (G) Wave area was measured after binarizing the denoised interpolated ratio image (upper). Box plots of the ERK wave area in each cell line at each frame from six to nine hours after releasing of silicone confinement. Data of three independent experiments are shown. p values of two-tailed t-test (4KO to others) are shown in panels B, C, D, F and G.

Next, we examined whether bath application of EGFR ligands could also restore the ERK activation waves. EGFR ligands at the same concentration, 10 ng mL^-1^ (∼ 2 nM), were added to 4KO cell lines after the removal of the confinement (Fig. S3A and Video 4). All EGFR ligands induced transient ERK activation and promoted leader cell migration (Fig. S3B and C). Nevertheless, none were able to induce ERK activation waves from the leader cells, indicating that only the endogenous EGFR ligands can generate ERK activation waves from the leader cells. Notably, the duration of the transient ERK activation was markedly longer in HBEGF-treated cells than in the cells treated with EGF, TGFα, or EREG (Fig. S3D).

### Growth arrest induced by the knockout of *Nrg1* in the absence of all four EGFR ligands

Albeit much less intensely than in WT cells, the ERK activation waves from the leader cells persisted even in QKO, 4KO, and dEGFR cells (Fig. 2A and Video 1), implying the involvement of neuregulins (NRGs), the ligands for ErbB3 and ErbB4. Among the four *Nrg* genes, only the expression of *Nrg1* was detected in MDCK cells (Fig. S1). Therefore, we attempted to knockout *Nrg1* in 4KO cells with three different sgRNAs targeted to exon 6 or exon 9 of *Nrg1*. Among 75 individual clones, 33 clones exhibited heterozygous deletion/insertion, but none showed homozygous knockout, suggesting that additional knockout of *Nrg1* in 4KO cells induces growth retardation. Therefore, we expressed *Nrg1* cDNA flanked by the *loxP* sequence before knockout of the endogenous *Nrg1* gene (Fig. 5A). This procedure successfully generated a clone named 5KO-loxP-NRG1, in which two-base deletion and single-base insertion were found in the *Nrg1* alleles (Fig. S2C). We further expressed Cre-ERT2 to generate 5KO-loxP-NRG1-CreERT2 cells and compared the cell growth in the presence and absence of 4-hydroxytamoxifen. As expected, after the addition of 4-hydroxytamoxifen, 5KO-loxP-NRG1-CreERT2 cells exhibited cell growth arrest (Fig. 5B), demonstrating the essential role of *Nrg1* in the growth of 4KO cells and indispensable role of the autocrine activation of the ErbB-family receptors in the growth of MDCK cells.

**Figure 5.**
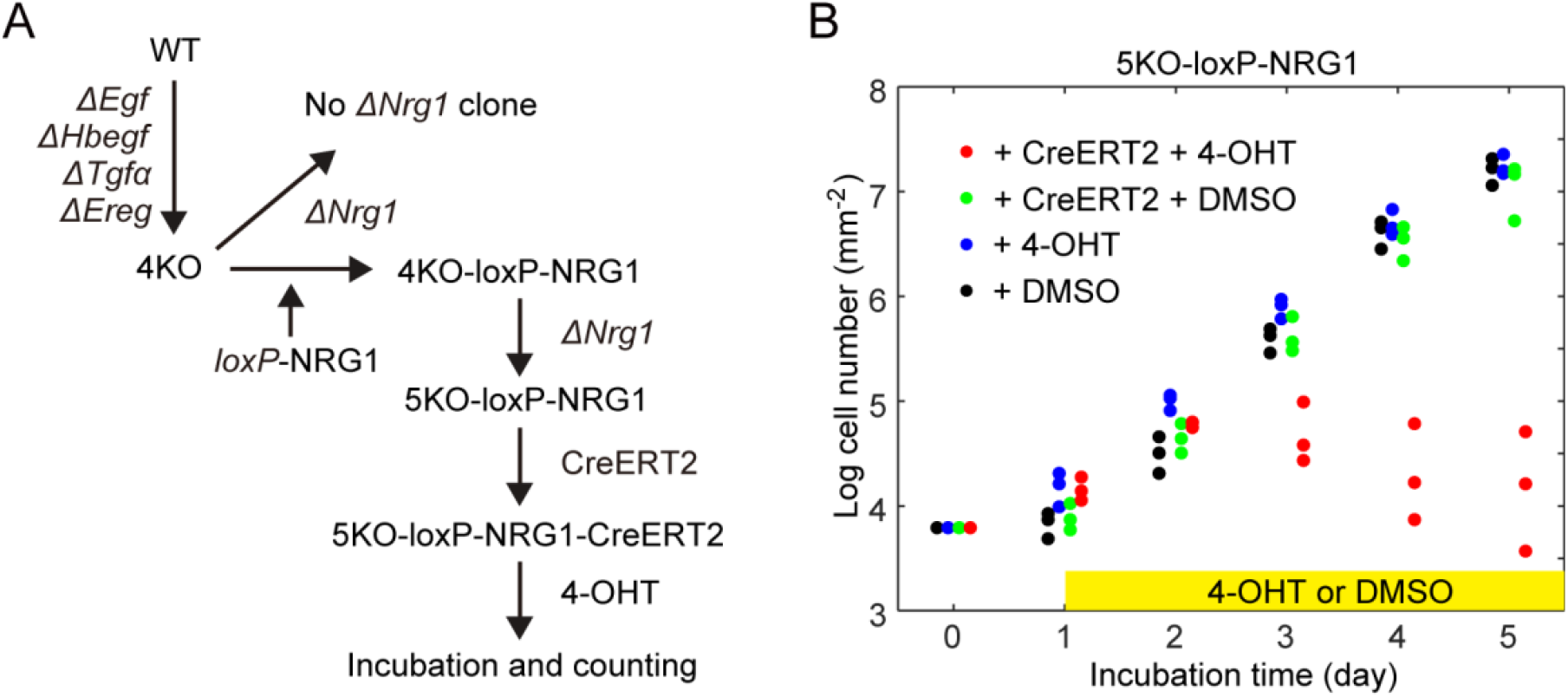
Growth arrest of cells deficient from all EGFR ligands. (A) Schematics of experiment to obtain 5KO cell line. The MDCK-4KO cells expressing EKARrEV biosensor were transfected stably with an expression vector for *loxP*-NRG1-*loxP* named as 4KO-loxP-NRG1. Then 5KO-loxP-NRG1 was generated by knocking out endogenous *Nrg1*. Followed by integrating CreERT2, 5KO-loxP-NRG1-CreERT2 was treated with 4-OHT to obtain 5KO cell line. (B) The growth rate represented on a log (cell number) basis. Cell numbers was measured over five days post-seeding of 5KO-loxP-NRG1 cell line with addition of 4-OHT (blue dots) or DMSO (black dots) and 5KO-loxP-NRG1-CreERT2 cell line with addition of 4-OHT (red dots) or DMSO (green dots) at day one. Each dot represents cell number in an independent experiment.

## Discussion

To examine the contribution of each EGFR ligand to the ERK activation waves, we first started to knockout the EGFR ligand genes from abundantly expressed *Egf* and *Hbegf*, followed by *Tgfα* and *Ereg*. Against our expectation, we found that the ERK activation waves from the leader cells were markedly inhibited only when all four EGFR ligands were knocked-out in QKO (Fig. 2E, Fig. 3C). A potential problem of our gene-knockout approach is that by this sequential knockout of the EGFR ligand genes, off-target mutations may accumulate. Therefore, we knocked-out *Hbegf, Egf, Tgfα* and *Ereg* simultaneously to obtain 4KO cells. By using these two independent clones, we minimized the potential effect of off-target mutation(s).

We previously reported that a disintegrin and metalloprotease 17 (ADAM17), also known as TACE, plays a critical role in the propagation of ERK activation waves in MDCK cells (Aoki et al., 2017; Hino et al., 2020). All pro-EGFR ligands except EGF are cleaved by ADAM17 to be soluble EGFR ligands (Sunnarborg et al., 2002). Then, why does EGF also restore the ERK activation waves? The ERK activation waves persist even in 4KO and QKO cells, suggesting the involvement of ErbB3 and ErbB4. EGF is the principal EGFR ligand that maintains basal ERK activity (Fig. 2), which is probably required for efficient ADAM17 activation (Fan and Derynck, 1999). Therefore, by restoring EGF-mediated ERK activation, NRG1, a substrate of ADAM17, may efficiently contribute to the ERK activation propagation in an ErbB3 and ErbB4-dependent manner. Another related unanswered question is that of why HBEGF failed to restore the ERK activation waves. Since the cell migration was accelerated by the HBEGF expression to the level of other EGFR ligands (Fig. 4B), functional HBEGF was expressed in 4KO-HBEGF cells. Because the auto-regulatory role of the heparin-binding domain of HBEGF may prevent proteolytic release in MDCK cells (Prince et al., 2010; Takazaki et al., 2004), ADAM17 activation caused by ERK activation waves may not be sufficient for the cleavage of HBEGF in MDCK cells. Alternatively, the prolonged ERK activation by HBEGF, for about 2 hours (Fig. S3D), may render HBEGF inappropriate for the propagation of the ERK activation waves, because duration of ERK activation is approximately a half hour both in MDCK cells (Fig. 2F) and in the mouse epidermis (Hiratsuka et al., 2015). Recently, in MCF10A human mammary epithelial cells, amphiregulin, expressed only at a marginal level in MDCK cells, was shown to mediate ERK activation from oncogene-expressing cells to neighboring normal cells (Aikin et al., 2020). Therefore, the dependency to each EGFR ligand and the redundancy appears cell type-specific.

The 4KO and QKO cells will provide a versatile platform to highlight the differences among EGFR ligands in a physiological context. Previously, extensive characterization of EGFR ligands has been conducted by bath application to tissue culture cells. There are at least two serious problems in this approach. First, bath application of EGFR ligands may not evoke the physiological phenotypes by autocrine stimulation. In fact, autocrine stimulation by EGFR ligands has been reported to promote greater cell migration of mammary epithelial cells compared to bath application (Joslin et al., 2007). In the present study, we also found that the ERK activation waves were restored by re-expression of EGFR ligands (Fig. 4A), but not by the bath application of these EGFR ligands (Fig. S3A). Second, the specific activity of each of the EGFR ligands used in previous studies might be significantly different. In one study using normal human epidermal keratinocytes, 2 nM EREG induced ERK phosphorylation more strongly than 10 nM EGF (Draper et al., 2003), whereas in another study using MCF-7 mammary cancer cells, 10 µM EREG and 16 nM EGF induced ERK phosphorylation to a similar level (Freed et al., 2017). In the present study, 10 ng mL^-1^ (∼ 2 nM) of EREG and EGF activated ERK to a similar level (Fig. S3B and Video 4). Although the cells used in each of these studies are different, these observations imply that the specific activity of EGFR ligands might be markedly different in each study. These potential flaws of bath application of EGFR ligands can be overcome by the re-expression of EGFR ligands in 4KO and QKO cells.

In conclusion, our results demonstrate that there is functional redundancy of EGFR ligands in the propagation of ERK activation waves during collective cell migration of MDCK cells. MDCK cells deficient in all EGFR ligands will provide a platform to examine the physiological function of each EGFR ligand.

## Materials and methods

### Cells, reagents, antibodies, plasmids, and primers

Reagents, antibodies, plasmids, and primers are described in the supplementary table and note.

### Cell culture

MDCK cells were from the RIKEN BioResource Center (no. RCB0995). Lenti-X 293T cells were purchased from Clontech (no. 632180; Mountain View, CA). These cells were maintained in DMEM (no. 044-29765; Wako, Osaka, Japan) supplemented with 10% FBS (no.172012-500ML; Sigma-Aldrich, St. Louis, MO), 100 unit mL^-1^ penicillin, and 100 mg mL^-1^ streptomycin (no. 26253-84; Nacalai Tesque, Kyoto, Japan) in a 5% CO2 humidified incubator at 37°C.

### cDNA cloning of dog EGFR ligands

Total RNA was isolated from MDCK cells using an RNeasy Mini Kit (no. 74104; Qiagen, Hilden, Germany) according to the manufacturer’s instructions. cDNA was reverse transcribed with a PrimeScript II 1st strand cDNA Synthesis Kit (no. 6210A; Takara Bio, Kyoto, Japan). Based on the information at RefSeq (http://www.ncbi.nlm.nih.gov/RefSeq), pairs of PCR primers specific to canine *Egf, Hbegf*, and *Ereg* were designed with an automated method utilizing Primer3 (https://primer3.org/) as listed in the supplementary table and note. The targeted ORF sequence was amplified by using KOD One PCR Master Mix (no. KMM-101; Toyobo, Osaka, Japan). The cDNA sequences were determined at the DNA Sequencing Facility of the Medical Research Support Center, Kyoto University. The cDNAs of canine *Tgfα* and *Nrg1* were synthesized by GeneArt (Thermo Fisher Scientific, Waltham, MA). The cDNA sequences of the cloned growth factors are shown in the supplementary table and note.

### Expression plasmids

The cDNAs of EGFR ligands were inserted into pPB-derived vectors (Yusa et al., 2009). CreERT2 cDNA (Matsuda and Cepko, 2007) was subcloned into pT2A-derived vector (Sumiyama et al., 2010) to generate pT2Aneo-CreERT2. For rapamycin-inducible activation of Ras, cDNA of Lyn-targeted FRB (LDR) and cDNA of mRFP-FKBP-mSos1-linkercat (Aoki et al., 2011) were subcloned into pPBpuro and pT2Aneo vector, respectively.

### CRISPR/Cas9-mediated KO cell lines

For CRISPR/Cas9-mediated single or multiple knockouts of genes encoding EGFR ligands or EGFR, single guide RNAs (sgRNA) targeting the exons were designed using CRISPRdirect (Naito et al., 2015). Oligo DNAs for the sgRNA were cloned into the lentiCRISPRv2 (Addgene Plasmid: no. 52961) vector or pX459 (Addgene Plasmid: no. 62988) vector. The expression plasmids for sgRNA and Cas9 were introduced into MDCK cells by lentiviral infection or electroporation. For lentivirus production, lentiCRISPRv2-derived expression plasmid, psPAX2 (Addgene Plasmid: no. 12260), and pCMV-VSV-G-RSV-Rev were co-transfected into Lenti-X 293T cells using polyethylenimine (no. 24765-1, Polyscience Inc., Warrington, PA). The infected cells were selected with media containing the following antibiotics, depending on the drug resistance genes carried by lentiCRISPRv2-derived plasmids; 100 µg mL^-1^ zeocin (no. 11006-33-0, InvivoGen, San Diego, U.S.A), 2.0 µg mL^-1^ puromycin (no. P-8833, Sigma-Aldrich, St. Louis, MO), 200 µg mL^-1^ hygromycin (no. 31282-04-9, Wako, Tokyo, Japan) and/or 800 µg mL^-1^ neomycin (no. 16512-52, Nacalai Tesque, Kyoto, Japan). For electroporation, pX459-derived expression plasmids were transfected into MDCK cells by an Amaxa Nucleofector II (Lonza, Basel, Switzerland). The transfected cells were selected with 2.0 µg mL^-1^ puromycin. After single cell cloning, genomic DNAs were isolated with SimplePrep reagent (no. 9180, TAKARA bio, Kyoto, Japan) according to the manufacturer’s instruction. PCR was performed using KOD FX neo (no. KFX-201 TOYOBO, Osaka, Japan) for amplification with designed primers, followed by DNA sequencing.

### Expression of FRET biosensors

cDNAs of EKARrEV-NLS were stably expressed either by lentivirus-mediated induction or transposon-mediated gene transfer. The pPB-derived vectors were co-transfected with pCMV-mPBase (neo-) (Yusa et al., 2009) at a ratio of 4:1 into MDCK cells using the Amaxa nucleofector system. Similarly, pT2A-EKAREV-NLS and pCAGGS-T2TP (Kawakami et al., 2004) were co-transfected into MDCK cells by electroporation. The established cell lines are summarized in the supplementary table and note.

### Time-lapse imaging by wide-field fluorescence microscopy

Fluorescence images were acquired essentially as described previously (Aoki and Matsuda, 2009). Briefly, cells cultured on glass-base dishes were observed under an IX83 inverted microscope (Olympus, Tokyo, Japan) equipped with a UPlanFL-PH 10x/0.3 (Olympus), a UPlanSApo 20x/0.75 (Olympus), or a UPlanSApo 40x/0.95 objective lens (Olympus), a DOC CAM-HR CCD camera (Molecular Devices, Sunnyvale, CA), a Spectra-X0 light engine (Lumencor Inc., Beaverton, OR), an IX3-ZDC laser-based autofocusing system (Olympus), an electric XY stage (Sigma Koki, Tokyo, Japan) and a stage top incubator (Tokai Hit, Fujinomiya, Japan). The filters and dichromatic mirrors used for time-lapse imaging were as follows: for FRET imaging, a 438/24 excitation filter incorporated in the Spectra-X light engine, a FF458-Di02-25×36 dichromatic mirror (Semrock, Rochester, NY), and FF01-483/32-25 and FF01-542/27-25 emission filters (Semrock) for CFP and FRET, respectively.

### Confinement release assay

The confinement release assay was performed as described previously (Hino et al., 2020). To observe collective cell migration of MDCK cells, a Culture-Insert 2 Well (no. 81176; ibidi, Martinsried, Germany) was placed on a 35 mm glass-base dish (no. 3911-035; IWAKI, Shizuoka, Japan) coated with 0.3 mg mL^-1^ type I collagen (Nitta Gelatin, Osaka, Japan). MDCK cells (3.5 × 10^4^) were then seeded in the Culture-Insert. Twenty-four hours after seeding, the silicone confinement was removed, and the medium was replaced with Medium 199 (11043023; Life Technologies, Carlsbad, CA) supplemented with 1% bovine serum albumin (BSA), 100 unit mL^-1^ penicillin, and 100 µg mL^-1^ streptomycin. Beginning at 30 min after the removal of the silicone confinement, the cells were imaged with an epifluorescence microscope every 2 or 5 min.

### Fluorescence lifetime imaging

For fluorescence lifetime imaging, 3.5 × 10^4^ MDCK cells were seeded in a Culture-Insert 2 well placed on a 24-well glass-bottom plate coated with 0.3 mg mL^-1^ type I collagen. Twenty-four hours after seeding, the silicone confinement was removed, and the medium was exchanged for Medium 199 supplemented with 1% BSA, 100 unit mL^- 1^ penicillin, and 100 µg mL^-1^ streptomycin. Six hours after the removal of the silicone confinement, the cells were imaged to measure the fluorescence lifetime of Turquoise. Lifetime imaging was performed with HC PL APO 20x/0.75 CS2 under a Leica TCS-SP8 microscope (Leica Microsystems GmbH, Wetzlar, Germany) equipped with a stage top incubator (Tokai Hit), a Lecia HyD SMD detector and a 440 nm picosecond pulsed diode laser (PDL 800-D; PicoQuant, Berlin, Germany), which pulsed at a frequency of 80 MHz. The band path of emission wavelength was set from 450 nm to 485 nm. Time-lapse images were acquired every 10 min. The acquisition time for each measurement was 45 seconds. The amplitude-weighted mean fluorescence lifetimes were calculated in a pixel-by-pixel fashion using fitting with a mono-exponential tail fit with adjustment of the number of components to two according to the manufacturer’s protocol (Leica Microsystems GmbH, Wetzlar, Germany).

### Western blotting with anti-phospho-ERK antibody

For the Western blotting analysis, 3.5 × 10^4^ cells MDCK cells were seeded in a single well of a Culture-Insert 2 Well that was placed on a glass-bottom dish coated with 0.3 mg mL^-1^ type I collagen. Twenty-four hours after seeding, the silicone confinement was removed. The medium was replaced with Medium 199 supplemented with 1% BSA, 100 unit mL^-1^ penicillin, and 100 µg mL^-1^ streptomycin. Six hours after the removal of the silicone confinement, MDCK cells were lysed with SDS sample buffer containing 62.5 mM Tris-HCl (pH 6.8), 12% glycerol, 2% SDS, 40 ng mL^-1^ bromophenol blue, and 5% 2-mercaptoethanol, followed by sonication with a Bioruptor UCD-200 (Cosmo Bio, Tokyo, Japan). After boiling at 95°C for 5 min, the samples were resolved by SDS-PAGE on SuperSep Ace 5-20% precast gels (Wako, Tokyo, Japan), and transferred to PVDF membranes (Merck Millipore, Billerica, MA) for Western blotting. All antibodies were diluted in Odyssey blocking buffer (LI-COR Biosciences, Lincoln, NE). Proteins were detected by an Odyssey Infrared Imaging System (LI-COR Biosciences, Lincoln, NE).

### Analysis of the cell migration distance in collective cell migration

To measure the migration distance of cells, the FIJI TrackMate plugin was applied to the CFP fluorescence images to acquire the track of each cell. The migration distance of each cell was defined by the difference between the abscissae of the first and last time points, twelve or eight hours later. The data analysis was performed by MATLAB.

### Quantification of ERK activity changes in single cells

Timelapse images of the FRET/CFP ratio were generated after background subtraction by using Metamorph software (Molecular Devices, Sunnyvale, CA) as described previously (Aoki and Matsuda, 2009). For single cell analysis of the ERK activity change, the images were analyzed by using the FIJI plug-in (Schindelin et al., 2012). On CFP images, the following commands were applied sequentially: “8-bit”, “;Subtract Background…” with the rolling size of 50 pixels, “Make Binary” with the Otsu method, “Watershed”, “Erode”, and “TrackMate” for tracking of each cell. The time-series data of the coordinates of each cell and the FRET/CFP ratio were processed with a Savitzky-Golay filter to reduce the noise.

The coefficient of duration *ω*_*i*_ and half of amplitude *A*_*i*_ were fitted as follows:

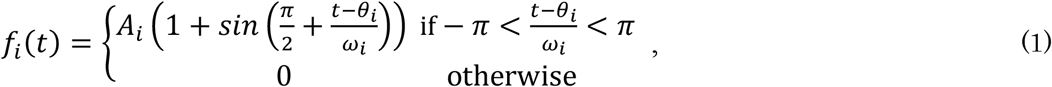

where *i* is the pulse index, t is the timepoint, and *θ*_*i*_ is the timepoint of the pulse peak. The programs used for sine curve fitting of MATLAB are shown in the Supporting Note.

### Analysis of ERK activation waves with heat maps

To determine the sample directions of ERK activation waves, heat maps of ERK activity were obtained by interpolating the signals in regions between the nuclei of MDCK cells in the FRET/CFP ratio images (Hino et al., 2020). The heat maps of ERK activity were analyzed by particle image velocimetry (PIV) using a free MATLAB-toolbox, MatPIV (Sveen, 2004), with a 128 µm window size and a 75% window overlap. The directions of the calculated velocity vectors were obtained as the sample directions. To obtain the kymographs of FRET/CFP ratios, these values were averaged along the y-axis in a defined region of the images, providing an intensity line along the x-axis. The operation was repeated for the respective time points, and the intensity lines were stacked along the y-axis for all time points. The ERK activation waves were detected and counted after binarizing by using the Regionprops (Image Processing Toolbox) function in MATLAB.

### Induction of ERK activation waves with rapamycin-inducible mSos1 translocation

The rapamycin-inducible mSos1 translocation and Ras activation were reported previously (Aoki et al., 2011). MDCK-WT-EKARrEV-NLS-LDR-mRFP-FKBP-mSos1-linkercat cells were seeded in a well of the Culture-Insert 1 well placed on a 24-well glass-bottom plate (no. 5826-024, IWAKI, Shizuoka, Japan). MDCK-WT-EKARrEV-NLS cells or MDCK-4KO-EKARrEV-NLS cells were seeded in the outside of the silicone confinement. After twenty-four hours incubation, the silicone confinement was removed. Further incubation for forty-eight hours allowed the cells to fill the gap between the two cell populations. During observation of the interface between the two cell populations, rapamycin was added to a final concentration of 250 nM. To examine the propagation of ERK activation waves, heat maps of ERK activity were obtained by interpolating the signals in regions between the nuclei of cells in the FRET/CFP ratio images.

### Analysis of the area of ERK activation waves

To examine the area of the ERK activation waves, heat maps of ERK activity were obtained by interpolating the signals in regions between the nuclei of cells in the FRET/CFP ratio images. To obtain a binarized image of the interpolated FRET/CFP ratio images, first these ratio images were denoised, and then a locally adaptive threshold was computed for a 2D grayscale image by using “adaptthresh” in MATLAB, after which the following arguments were applied sequentially: “sensitivity” of 0.65, and “ForegroundPolarity” with “dark”. After obtaining binarized images, wave areas of each frame in each cell line from six to nine hours after releasing the silicone confinement was counted.

### Analysis of the duration of the transient ERK activation after addition of recombinant human EGFR ligands

To examine the duration of the transient ERK activation after application of recombinant human EGFR ligands, time of ERK activation recovering to the half-maximal were calculated. The definition of half-maximal is the average of ratio value before EGFR ligands adding (basal) and just after adding (maximum) in each experiment.

### Measurement of growth rate

For the growth rate measurement, 5 × 10^4^ MDCK-5KO-EKARrEV-NLS-loxP-NRG1 or MDCK-5KO-EKARrEV-NLS-loxP-NRG1-CreERT2 cells were seeded in a 6-well plate (No. 140675, Thermo Fisher Scientific, Waltham, MA) and cultured in DMEM containing 10% FBS. After twenty-four hours incubation, 1 µM 4-hydroxytamoxifen (no. 579002, Sigma-Aldrich, St. Louis, MO) or DMSO (no. 13445-74, Nacalai Tesque, Kyoto, Japan) was added. Fluorescence images of fixed locations were acquired with a UPlanFL-PH 10x/0.3 objective lens (Olympus) every twenty-four hours. Images were binarized to count the number of nuclei by FIJI.

### Characterization of EKARrEV-NLS

We first established MDCK cells stably expressing EKARrEV-NLS or the prototype EKAREV-NLS. Then, 2 × 10^4^ cells were seeded in a well of a 24-well glass-bottom plate coated with 0.3 mg mL^-1^ type I collagen. After twenty-four hours, the medium was replaced with Medium 199 supplemented with 1% BSA, 100 unit mL^-1^ penicillin, and 100 µg mL^-1^ streptomycin. Fluorescence images were acquired with an UPLSAPO 20X objective (Olympus) every 2 min. During imaging, EGF (no. E9644; Sigma-Aldrich, St. Louis, MO) or trametinib (no. T-8123; LC Laboratories, Woburn, MA) was added to a concentration of 100 ng mL^-1^ or 200 nM final, respectively. After applying autothreshold, the fluorescence intensities of each nucleus were quantified to obtain FRET/CFP values as described previously (Aoki and Matsuda, 2009).

### Statistical analysis

Probability (p) values were determined by using the T.TEST function of Microsoft Excel with two-tailed distribution and two-sample unequal variance.

## Acknowledgements

We thank the members of the Matsuda Laboratory for their helpful input and encouragement, K. Hirano and K. Takakura for their technical assistance, Takahiko Matsuda for pCAG-CreERT2, and the Medical Research Support Center of Kyoto University for in vivo imaging. This work was supported by the Kyoto University Live Imaging Center. Financial support was provided in the form of JSPS KAKENHI grants (nos. 18K07066 to K.T., 20H05898 to M.M., and 19H00993 to M.M.), a JST CREST grant (no. JPMJCR1654), a Moonshot R&D grant (no. JPMJPS2022 to M.M.), and funds from the Fugaku Foundation (to M.M.).

## Author contributions

S.L., K.A., N.H., K.T., and M.M. conceived and designed the study. S.L., D.H., G.M., K.M., N.H., and E.D. acquired and analyzed the experimental data. R.I. provided plasmids and cells. S.L. and M.M. wrote the manuscript.

---

**Video 1. ERK activation waves in WT and mutant MDCK cells during collective cell migration, related to figure 2A**

Time-lapse movie of collectively migrating MDCK cells expressing EKARrEV-NLS. The golden pseudo-color represents the FRET/CFP ratio indicating ERK activity. The color scales correspond to those in figure 2A. Time in h:min.

**Video 2. Propagation of ERK activation waves upon rapamycin induced ERK activation, related to figure 3E**

Time-lapse movie of the boundary between confluent MDCK WT (left) or 4KO (right) cells with rapamycin-activatable mSos1 expression WT cells. The color represents the FRET/CFP ratio indicating ERK activity. The cells were treated with 250 nM rapamycin at 0 min to induce ERK activation of the RA mSos1 expressing cells. Time in h:min.

**Video 3. ERK activation waves in WT, 4KO and 4KO expressing EGFR ligands MDCK cells during collective cell migration, related to figure 4A**

Time-lapse movie of collectively migrating MDCK cells expressing EKARrEV-NLS. The golden pseudo-color represents the FRET/CFP ratio indicating ERK activity. The color scales correspond to those in figure 4A. Time in h:min.

**Video 4. Restored leader cell migration upon addition of recombinant EGFR ligands, related to figure S3A**

Time-lapse movie of collectively migrating MDCK cells expressing EKARrEV-NLS. The golden pseudo-color represents the FRET/CFP ratio indicating ERK activity.

MDCK 4KO cells were treated with 10ng mL^-1^ recombinant human EGF, HBEGF, TGFα or EREG at 0 min. Time in h:min.

## Supplementary figures

**Figure S1.**
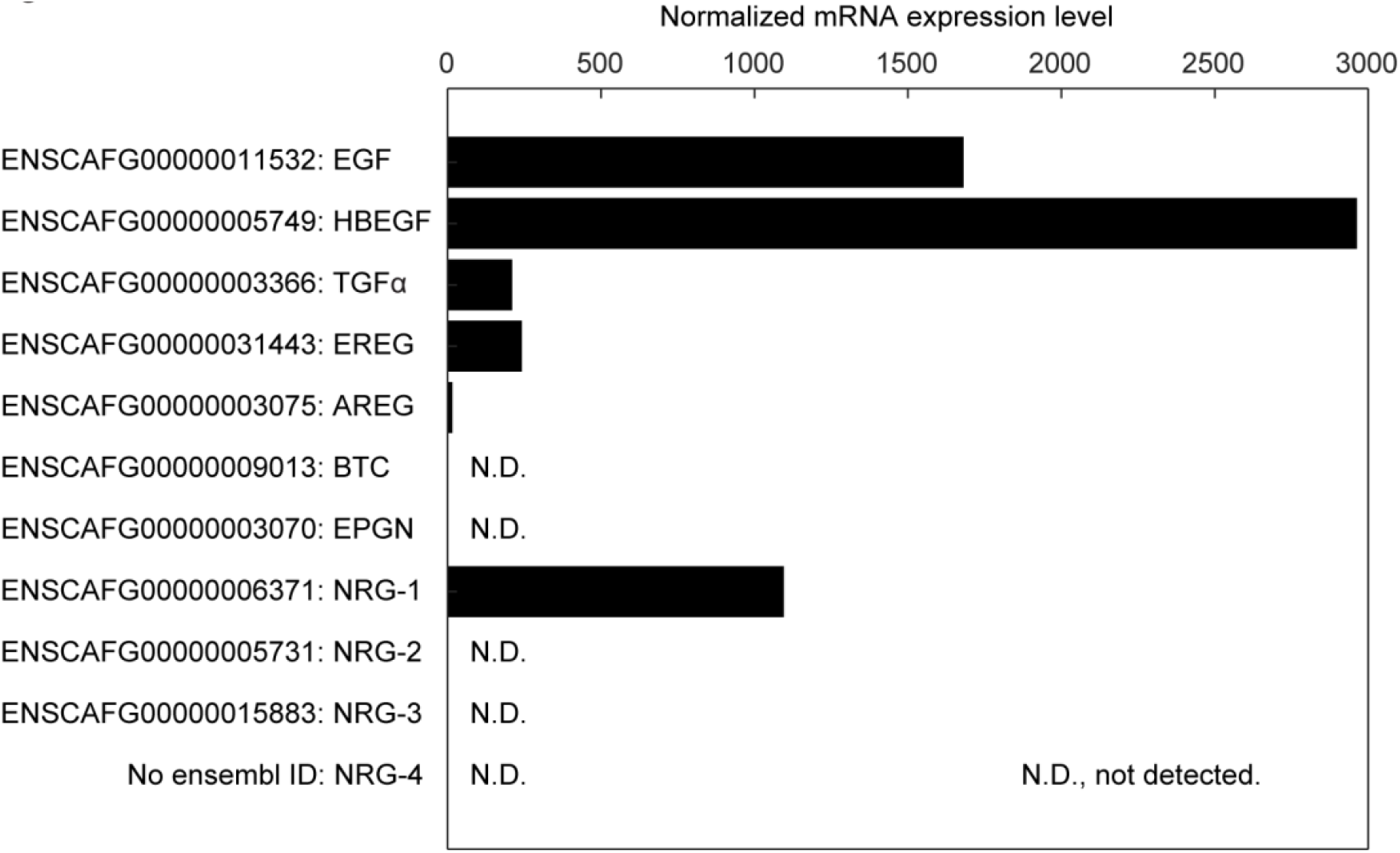
mRNA expression level of EGFR ligands in MDCK cells. Normalized mRNA expression level of EGFR ligands in MDCK cells calculated from RNA-Seq data (Shukla et al., 2015). N.D., not detected.

**Figure S2.**
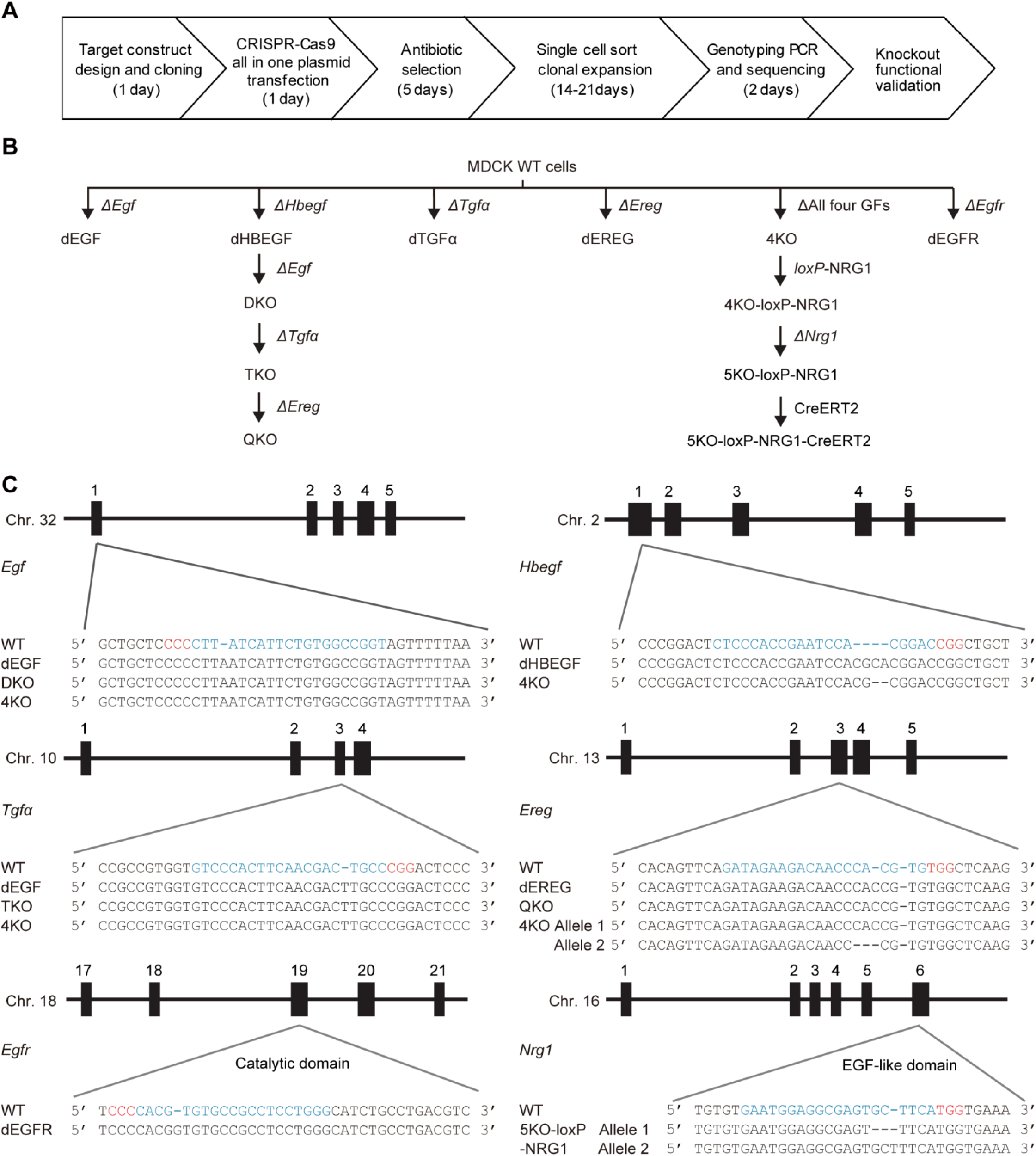
Generation of gene knockouts. (A) Workflow for generation of gene knockouts in MDCK cell lines. (B) Phylogenetic tree of the MDCK cells used in this study. (C) Mutations of the targeted genes in the MDCK cell lines. Black rectangles depict exons. The gRNA and PAM sequences are depicted in blue and red, respectively.

**Figure S3.**
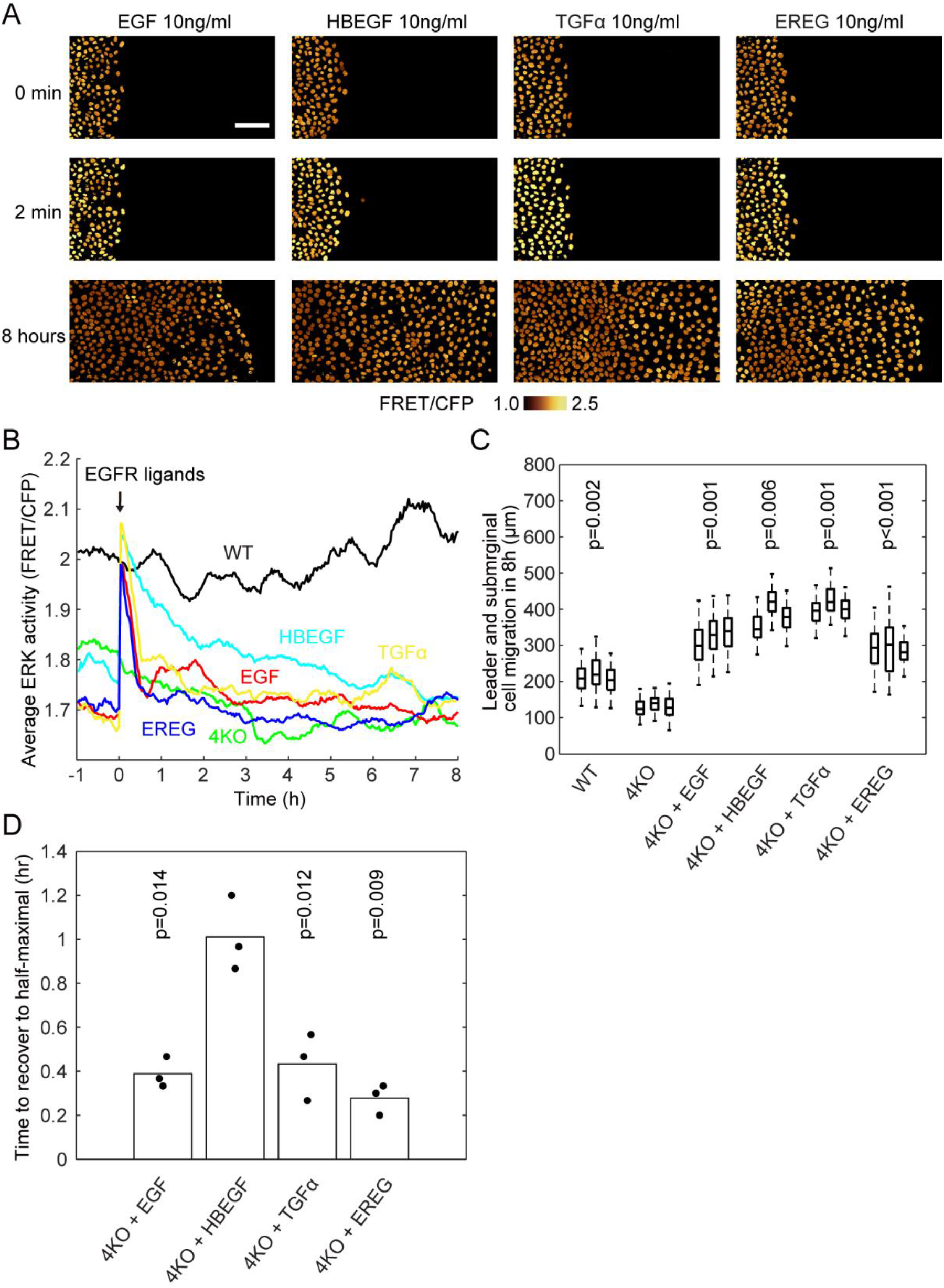
Restoration of migration by addition of EGFR ligands in 4KO cells. (A) 4KO cells were subjected to confinement release assay as described in the legend to Figure 1. Images of CFP and FRET channels were acquired every 2 minutes for twelve hours (Video 4). Shown are snapshots before, just and 8 hours after the addition of 10 ng mL^-1^ recombinant EGFR ligands. Scale bar, 100 μm.(B) Time course of the average FRET/CFP ratio. At time 0 hour, 10ng/ml recombinant human EGF (red), HBEGF (light blue), TGFα (yellow) and EREG (blue) were added to 4KO cell lines. (C) Leader cell migration in eight hours after the addition of EGFR ligands or after the release of silicone confinement (WT and 4KO). The numbers of analyzed cells are as follows: WT, 357, 215 and 283 cells; 4KO, 348, 274, and 328 cells; 4KO + EGF, 154, 128, and 193 cells; 4KO + HBEGF, 160, 253, and 235 cells; 4KO + TGFα, 301, 274, and 213 cells; 4KO + EREG, 236, 219, and 86 cells. Data of three independent experiments are shown. p values of two-tailed t-test (4KO to others) are shown in the panel. (D) Time to the half-maximal activity after the addition of EGFR ligands. Data of three independent experiments are shown. p values of two-tailed t-test (4KO + HBEGF to others) are shown in the panel.

