## supplementary table and note for "Redundant and specific roles of EGFR ligands in the ERK activation waves during collective cell migration of MDCK cells"

### Reagents

| Reagent | Source | Identifier |
| --- | --- | --- |
| Dimethyl Sulfoxide | Nacalai Tescue | 13445-74 |
| Trametinib | LC Laboratories | T-8123 |
| Rapamycin | LC Laboratories | R-5000 |
| Zeocin | InvivoGen | 11006-33-0 |
| Hygromycin | Wako | 31282-04-9 |
| Neomycin | Nacalai Tescue | 16512-52 |
| Penicillin and Streptomycin | Nacalai Tescue | 26253-84 |
| 4-Hydroxytamoxifen | Sigma-Aldrich | 579002 |
| Recombinant Human EGF | Sigma-Aldrich | E9644 |
| Recombinant Human HBEGF | PeproTech | 100-47 |
| Recombinant Human TGFα | R&D | 239-A-100 |
| Recombinant Human EREG | PeproTech | 100-04 |
| Bovine Serum Albumin | Sigma-Aldrich | A2153 |
| Fetal bovine serum | Sigma-Aldrich | 172012-500ML |

### Antibody

| Antibody | Source | Identifier |
| --- | --- | --- |
| Anti-ERK1/2 mouse antibody | BD Biosciences | Cat# 610123, RRID:AB_397529 |
| IRDye 680-conjugated goat anti-mouse IgG antibody | LI-COR Biosciences | Cat# 926-32220, RRID:AB_621840 |
| IRDye 800CW goat anti-rabbit IgG antibody | LI-COR Biosciences | Cat# 926-32211, RRID:AB_621843 |
| Anti-phospho-p44/42 MAPK (Erk1/2; Thr202/Tyr204) rabbit antibody | Cell Signaling Technology | Cat# 9101, RRID:AB_331646 |

### Plasmid

| Plasmid | Source | Identifier |
| --- | --- | --- |
| pCAGGS-T2TP | a gift from Koichi Kawakami (National Institute of Genetics, Japan) | N/A |
| pCX4puro-LDR | (Li et al, 2017) | N/A |
| pCMV-VSVG-RSV-Rev | (Miyoshi et al., 1998) | N/A |
| psPAX2 | a gift from Didier Trono | Addgene plasmid # 12260 |
| pLentiCRISPRv2 | This manuscript | Addgene plasmid # 52961 |
| pT2A-EKAREV-NLS | (Kawabata et al., 2016) | N/A |
| pPBbsr-EKARrEV-NLS | This manuscript | N/A |
| pPBpuro-UbC-MCS | a gift from Dr. Kosuke Yusa, Team 83, Welcome Trust Sanger Institute | N/A |
| pCMV-mPBase | a gift from Dr. Kosuke Yusa, Team 83, Welcome Trust Sanger Institute | N/A |
| pCAG-CreERT2 | a gift from Dr. Takahiko Matsuda | N/A |
| pX459 (pSpCas9(BB)-2A) | a gift from Dr. Feng Zhang | Addgene plasmid # 62988 |
| pMcsbsr-mRFP-FKBP-mSos1-linkercat | (Aoki et al., 2011) | N/A |
| pT2Aneo-CreERT2 | This manuscript | N/A |
| pPBpuro-LDR | This manuscript | N/A |
| pT2Aneo-mRFP-FKBP-mSos1-linkercat | This manuscript | N/A |
| pPBpuro-UbC-loxP-NRG1 | This manuscript | N/A |

### Origianl cell lines

| Name | Source | Identifier |
| --- | --- | --- |
| Dog: MDCK cells | RIKEN BioResource Center | RCB0995 |
| Human: Lenti-X 293T cells | Clontech | 632180 |

### Established cell lines

| Name | Short name | Original cell line | Plasmid |
| --- | --- | --- | --- |
| MDCK-EKARrEV-NLS | WT | MDCK-WT | pCSIIbsr-EKARrEV-NLS, pCMV-VSV-G-RSV-Rev, psPAX2 |
| MDCK-EKAVEV-NLS | N/A | MDCK-WT | pT2A-EKAVEV-NLS, pCAGGS-T2TP |
| MDCK-dEGF | N/A | MDCK-WT | pLentiCRISPRv2, pCMV-VSV-G-RSV-Rev, psPAX2 |
| MDCK-dHBEGF | N/A | MDCK-WT | pLentiCRISPRv2, pCMV-VSV-G-RSV-Rev, psPAX2 |
| MDCK-dTGFα | N/A | MDCK-WT | pLentiCRISPRv2, pCMV-VSV-G-RSV-Rev, psPAX2 |
| MDCK-DKO | N/A | MDCK-dHBEGF | pLentiCRISPRv2, pCMV-VSV-G-RSV-Rev, psPAX2 |
| MDCK-TKO | N/A | MDCK-DKO | pLentiCRISPRv2, pCMV-VSV-G-RSV-Rev, psPAX2 |
| MDCK-QKO | N/A | MDCK-TKO | pLentiCRISPRv2, pCMV-VSV-G-RSV-Rev, psPAX2 |
| MDCK-4KO | N/A | MDCK-WT | pX459 |
| MDCK-dEGF-EKARrEV-NLS | dEGF | MDCK-dEGF | pCSIIbsr-EKARrEV-NLS, pCMV-VSV-G-RSV-Rev, psPAX2 |
| MDCK-dHBEGF-EKARrEV-NLS | dHBEGF | MDCK-dHEBGF | pCSIIbsr-EKARrEV-NLS, pCMV-VSV-G-RSV-Rev, psPAX2 |
| MDCK-dTGFα-EKARrEV-NLS | dTGFα | MDCK-dTGFα | pCSIIbsr-EKARrEV-NLS, pCMV-VSV-G-RSV-Rev, psPAX2 |
| MDCK-EKARrEV-NLS-dEREG | dEREG | MDCK-EKAVrEV-NLS | pX459 |
| MDCK-DKO-EKARrEV-NLS | DKO | MDCK-DKO | pCSIIbsr-EKARrEV-NLS, pCMV-VSV-G-RSV-Rev, psPAX2 |
| MDCK-TKO-EKARrEV-NLS | TKO | MDCK-TKO | pCSIIbsr-EKARrEV-NLS, pCMV-VSV-G-RSV-Rev, psPAX2 |
| MDCK-QKO-EKARrEV-NLS | QKO | MDCK-QKO | pCSIIbsr-EKARrEV-NLS, pCMV-VSV-G-RSV-Rev, psPAX2 |
| MDCK-4KO-EKARrEV-NLS | 4KO | MDCK-4KO | pPBbsr2-EKARrEV-NLS |
| MDCK-EKARrEV-NLS- dEGFR | dEGFR | MDCK-EKAVrEV-NLS | pLentiCRISPRv2, pCMV-VSV-G-RSV-Rev, psPAX2 |
| MDCK-WT-EKARrEV-NLS-LDR | N/A | MDCK-WT-EKARrEV-NLS | pPBpuro-LDR, pCMV-mPBase(neo-) |
| MDCK-WT-EKARrEV-NLS-LDR-mRFP-FKBP-mSos1-linkercat | RA-mSos1 | MDCK-WT-EKARrEV-NLS-LDR | pT2Aneo-mRFP-FKBP-mSos1-linkercat, pCAGGS-T2TP |
| MDCK-4KO-EKARrEV-NLS-EGF | 4KO-EGF | MDCK-4KO-EKARrEV-NLS | pPB-UbC-canisEGF, pCMV-mPBase(neo-) |
| MDCK-4KO-EKARrEV-NLS-HBEGF | 4KO-HBEGF | MDCK-4KO-EKARrEV-NLS | pPBpuro-UbC-canisHBEGF, pCMV-mPBase(neo-) |
| MDCK-4KO-EKARrEV-NLS-TGFα | 4KO-TGFα | MDCK-4KO-EKARrEV-NLS | pPB-UbC-canisTGFα, pCMV-mPBase(neo-) |
| MDCK-4KO-EKARrEV-NLS-EREG | 4KO-EREG | MDCK-4KO-EKARrEV-NLS | pPBpuro-UbC-canisEREG, pCMV-mPBase(neo-) |
| MDCK-4KO-EKARrEV-NLS-loxP-NRG1 | 4KO-loxP-NRG1 | MDCK-4KO-EKARrEV-NLS | pPBpuro-UbC-loxP-canisNRG1, pCMV-mPBase(neo-) |
| MDCK-5KO-EKARrEV-NLS-loxP-NRG1 | N/A | MDCK-4KO-EKARrEV-NLS-loxP-NRG1 | pX459 |
| MDCK-5KO-EKARrEV-NLS-loxP-NRG1-CreERT2 | 5KO-loxP-NRG1-CreERT2 | MDCK-5KO-EKARrEV-NLS-loxP-NRG1 | pT2Aneo-CreERT2, pCAGGS-T2TP |

### Primers

| Primers | Details |
| --- | --- |
| Canine EGF clone Fw | 5’- AGCAGAATTCATGCTGCTCCCCCTTATCA -3’ |
| Canine EGF clone Rv | 5’- TTAGCGGCCGCTCACTGAATCAGCTTCA -3’ |
| Canine HBEGF clone Fw | 5’- AGCAGAATTCATGAAGCTGCTGCGGTCAG -3’ |
| Canine HBEGF clone Rv | 5’- TTAGCGGCCGCCTAGCTGTGACGTCCTT -3’ |
| Canine EREG clone Fw | 5’- ATTAGAATTCATGGAGCCGCGCCGCCTGC -3’ |
| Canine EREG clone Rv | 5’- TTAGCGGCCGCTCAGACTTGTGGCAATG -3’ |
| sgRNA for canine EGF | 5’- CTTATCATTCTGTGGCCGGT -3’ |
| sgRNA for canine HBEGF | 5’- CTCCCACCGAATCCACGGAC -3’ |
| sgRNA for canine TGFα | 5’- GTCCCACTTCAACGACTGCC -3’ |
| sgRNA for canine EREG | 5’- GATAGAAGACAACCCACGTG -3’ |
| sgRNA for canine NRG1 | 5’- GAATGGAGGCGAGTGCTTCA -3’ |
| sgRNA for canine EGFR | 5’- CACGTGTGCCGCCTCCTGGG -3’ |
| Detecting canine EGF Fw | 5’- CTGAACGTCCGTTTCAGGCT -3’ |
| Detecting canine EGF Rv | 5’- ACACTCCGAGAGAAATCAGGC -3’ |
| Detecting canine HBEGF Fw | 5’- GGGTGCAGAGGGGTCTGATG -3’ |
| Detecting canine HBEGF Rv | 5’- GCAAGCTTGTCGGGTTTAGC -3’ |
| Detecting canine TGFα Fw | 5’- TGCAAAAGTTAAGGTGCGGC -3’ |
| Detecting canine TGFα Rv | 5’- TTTGAAGCAGGTGTCGCTCA -3’ |
| Detecting canine EREG Fw | 5’- CCTTAGGGGTCAAATGTTTGTTTAGTGGTG -3’ |
| Detecting canine EREG Rv | 5’- GCTTTGTAAGTTTTGTAGGTTTGAGCTGCC -3’ |
| Detecting canine NRG1 Fw | 5’- CCTCCGTACCTTATGTGAACTC -3’ |
| Detecting canine NRG1 Rv | 5’- CTGGATGTAACAGAGGAGATGA -3’ |
| Detecting canine EGFR Fw | 5’- TGCCTGGGTGTCTCTGTGAAGGGAAATCCT -3’ |
| Detecting canine EGFR Rv | 5’- CCCACCTGTCCACTTTCCGGCTGTCTTTAT -3’ |

### EKARrEV-NLS

YPet, WW domain, EV linker, ERK substrate peptide, mTurquoise, NLS

ATGGTGAGCAAGGGCGAGGAGCTGTTCACCGGGGTGGTGCCCATCCTGGTCGAGCTGGACGGCGACGTAAACGGCCACAAGTTCAGCGTGTCCGGCGAGGGCGAGGGCGATGCCACCTACGGCAAGCTGACCCTGAAGCTTCTATGCACCACCGGCAAGCTGCCCGTGCCCTGGCCCACCCTCGTGACCACCCTGGGCTACGGCCTGCAGTGCTTCGCCCGCTACCCCGACCACATGAAGCAGCACGACTTCTTCAAGTCCGCCATGCCCGAAGGCTACGTCCAGGAGCGCACCATCTTCTTCAAGGACGACGGCAACTACAAGACCCGCGCCGAGGTGAAGTTCGAGGGCGACACCCTGGTGAACCGCATCGAGCTGAAGGGCATCGACTTCAAGGAGGACGGCAACATCCTGGGGCACAAGCTGGAGTACAACTACAACAGCCACAACGTCTATATCACCGCCGACAAGCAGAAGAACGGCATCAAGGCCAACTTCAAGATCCGCCACAACATCGAGGACGGCGGCGTGCAGCTCGCCGACCACTACCAGCAGAACACCCCCATCGGCGACGGCCCCGTGCTGCTGCCCGACAACCACTACCTGAGCTACCAGTCCGCCCTGTTCAAAGACCCCAACGAGAAGCGCGATCACATGGTCCTGCTGGAGTTCCTGACCGCCGCCGGGATCACTGAGGGCATGAACGAGCTGTACCTCGAGATGGCGGACGAGGAGAAGCTGCCGCCCGGCTGGGAGAAGCGCATGAGCCGCAGCTCAGGCCGAGTGTACTACTTCAACCACATCACTAACGCCAGCCAGTGGGAGCGGCCCAGCGGCAACAGCAGCAGTGGTGGCAAAAACGGGCAGGGGGAGCCTGCCAGGGGTACCAGTGCTGGTGGTAGTGCTGGTGGTAGTGCTGGTGGTAGTGCTGGTGGTAGTGCTGGTGGTTCCGGCAGTGCTGGTGGTAGTGCTGGTGGTAGTACCAGTGCTGGTGGTAGTGCTGGTGGTAGTGCTGGTGGTAGTGCTGGTGGTAGTGCTGGTGGTTCCGGCAGTGCTGGTGGTAGTGCTGGTGGTAGTACCAGTGCTGGTGGTAGTGCTGGTGGTAGTGCTGGTGGTAGTGCTGGTGGTAGTGCTGGTGGTTCCGGCAGTGCTGGTGGTAGTGCTGGTGGTAGTACCAGTGCTGGTGGTAGTGCTGGTGGTAGTGCTGGTGGTAGTGCTGGTGGTAGTGCTGGTGGTTTCCGGACTCCTGATGACGCCATGCTACACGGCAAAGCTGTCATTCCAATTTCCGGGCGGCCGCATGGTGAGCAAGGGCGAGGAGCTGTTCACCGGGGTGGTGCCCATCCTGGTCGAGCTGGACGGCGACGTAAACGGCCACAGGTTCAGCGTGTCCGGCGAGGGCGAGGGCGATGCCACCTACGGCAAGCTGACCCTGAAGTTCATCTGCACCACCGGCAAGCTGCCCGTGCCCTGGCCCACCCTCGTGACCACCCTGACCTGGGGCGTGCAGTGCTTCAGCCGCTACCCCGACCACATGAAGCAGCACGACTTCTTCAAGTCCGCCATGCCCGAAGGCTACGTCCAGGAGCGTACCATCTTCTTCAAGGACGACGGCAACTACAAGACCCGCGCCGAGGTGAAGTTCGAGGGCGACACCCTGGTGAACCGCATCGAGCTGAAGGGCATCGGCTTCAAGGAGGACGGCAACATCCTGGGGCACAAGCTAGAGTACAACTACATCAGCCACAACGTCTATATCACCGCCGACAAGCAGAAGAACGGCATCAAGGCCCACTTCAAGATCCGCCACAACATCGAGGACGGCGGCGTGCAGCTCGCCGACCACTACCAGCAGAACACCCCCATCGGCGACGGCCCCGTGCTGCTGCCCGACAACCACTACCTGAGCACCCAGTCCGCCCTGAGCAAAGACCCCAACGAGAAGCGCGATCACATGGTCCTGCTGGAGTTCGTGACCGCCGCCGGGATCACTCTCGGCATGGACGAGCTGTCTAGACCTAAGAAGAAGCGTAAGGTG

### Canine EGF

ATGCTGCTCCCCCTTATCATTCTGTGGCCGGTAGTTTTTAAATGTAGTTTTGCTAGTCTCTCAGACCCGGAGAACTGGAACTGTCCTGAAGTCTCTCCCTCAGGAAAGGGGAGCCCTGCTTGTGTGGGTCCTGCACCCTTCTTAATTTTCTCCCATGGAATCAGTATCTTTAGGATTGACCTGGAAGGCACTAATCATGAGCAATTGGTGGCAGATGCTGGTGTATCAGTGATCATGGATTTTCATTATAATAAAGAAAGAATCTATTGGGTAGATCCAGAAAGACAACTTTTGCAAAGAGTTTTTCTGAATGGGACAAGGCAAGAGAGAGTATGCAATATAGAGAAAAATGTTTCAGGAATGGCAATAAATTGGATAAATGAAGAACTTATTTGGTCAAATCAACAGGAAGGAATCATCACAGTAACAGATATGAAAGGAAACAATTCGCGAGTTCTCTTAAGAGCCTTAAACTATCCTGCAAATGTAGCAATTGATCCAATAGAAAGGTTTATATTTTGGTCTTCAGAGGTGGCAGTGGCTGGCAGCCTTCACAGAGCAGATCTCAATGGTGTGGAAGAGAAGATTCTTTTACAGACATCAGAAAGAATAACAGCTGTGTCATTGGATGTGCTTGATAAACAGCTTTTTTGGATTCAGTACAGCAGAGATGGAAGCAATTCTCATATTTATTCCTGTAATTATGATGGAGGTTCTGTCCATCTTAGCAAACATCTTACACAGCATAATTTTTTTGCAATGTCCCTTTTTGGGAATCAGATCTTCTATTCAACATGGAAAAAGAAGACAATTTGGATAGCTAACAAACACAGTGGGAAGGATATGGTTAGAATTAACCTAGATTCATCATTTGTACCACCTGGTGGAATCAAAGTAGTGCATCCACTCTTACAGCCCAAGGCAGAGAGTGGCACTTGGGCACCTGATCAGAAACTCTGCAAATGGAAGCAAGGTAACTGCAGAGGCAGCACGTGTGGTCAAGATTCAAAGTCCTATTCATGCACGTGTGCAGAGGGATATACTTTAAGCCAAGATGGAAAATACTGTGAAGATGTCAATGAATGTGCCTTTTGGAATCATGGCTGTACTCTTGGGTGTGAAAACATCCCTGGATCTTATTATTGCACATGCCCTGTAGGATTTATTCTGCTTCCTGATGGGAAACGGTGTCATCAATTAATTGCCTGTCCGAGCAATACATCTAAATGTAGCCATGACTGTGTTCTGACATCAGATGGTCCCATATGTTTCTGTCCTGAAGGCTCAGTGCTTGAGGCAGATGGAAAAACATGTAGTGGCTGTTCATCACCTGATAATGGTGGATGTAGCCAGCTCTGCCTCCCTCTCAGCCCAGTATCCTGGGAATGTGGTTGCTTTCCTGGGTATGACCTACAACTGGACAAACAAAGCTGTGCAGCATCAGGACCACAACCATTTTTGCTGTTTGCCAATTCTCAAGATATTCGACACATGCATTTTGATGGAACAGATTATGGAACCCTGCTCAGCCAGCAAATGGGAATGGTTTTTGCCCTTGATCACGACCCTGTGGAAAATAAGATATACTTTGCCCATACAGCCCTGAAGTGGATAGAGAGAGCTAACATGGATGGTTCCCAGCGAGAAAGGCTTATTGAGGAAGGAGTGGATGTGCCAGAAGGTCTTGCCATAGATTGGATTGACCGTAAATTCTACTGGACAGACAGCGGAAAATCTCTTATTGAAGGGAGTGATTTAAATGGAAAACATCGTGAGGTAATCATCAAGGAAGACATCTCTCAGCCACGAGGAATTGCTGTTCATCCAATGGCCAAGAGATTATTCTGGACTGATATGGGGATTAATCCACGAATTGAAAGTTCTTCCCTTCAAGGCATTGGCCGACTGGTTATAGCTAGCTCTGATCTGGTCTGGCCCAGTGGAATAACGATTGATTATGTAACTGACAAATTGTACTGGTGTGATACCAAGCTGTCTGTGATTGAGATGGCCAATCTGGATGGTTCAAAACGCCAAAGACTTGCCCAGAACGATGTAGGTCACCCATTTGCTATGGCCGTGTTTGAGGATCACGTGTGGTTCTCTGATTGGACTATGCCATCAATAATAAGAGTGGACAAGAGGACTGGCAAAAACAGGGTACGTCTCCGAGGCAGCATGCTGAAGCCTTCATCACTGGTTGTAGTTCATCCATTGGCAAAACCAGGAGCACAGCCCTGCTTATATCAAAATGGAGGCTGTGAACATATTTGCAAAGAGAGGTTTGGAACTGCTCAATGTTTGTGTCGTGAAGGTTTTGTGAAAGCCCCAGATGGGAAAATGTGTCTGGCTCTGAATGGCCATCAGATACCGGCAGTAGGTAGTGAAGCAGATCTAAGTAATCACGTAACGCCAGGGGATGTCTTACCCAGAAGTGAAGGATTTGAAGATAACATTACAGAATCTCAGCATATGCTAGTGGCCGAAATCATGGTGTCAGATGCTGACGACTGTGCTCCTGTGGGATGCAGTACATGGGCTGAGTGTGTTTCAGAGGGAGAAAATGCCACATGTCAGTGTTTGAAAGGATTTACTGGGGATGGAAAGCTATGTTTTGACATAGATGAATGTGAGATGGGCATCACGATTTGCCCTCCTACCTCCTCAAAGTGCGTCAATACTGAAGGTGGTTATGTTTGTCAGTGCTCAGAAGGCTACCGAGGCGATGGGATCCACTGTCTGGATATTAATGAGTGCCAACTGGGCATGCACACCTGTGGGGAAAATGCCACCTGTACAAATATGGAGGGAAACTATACCTGCATGTGCGCTGGCAGCCTGTCTGAACCTGGACAGATATGTGCTGACTCTACTCCGCCTTCTCATCCCATGGAGGACAGTCACTATTCTGTGAGAAATGGTTATCGGGAATGCCCCTCATCCTATGATGGGTACTGCCTCTATAATGGTGTGTGTATGTACATTGAAGCAGTCGACAGATACGCATGCAACTGTGTTTTTGGCTACGTCGGGGAGCGATGTCAGCACCGAGACCTGAAATGGGAACTGCGCCACGCGGGCCAGGGCCGGCAGCGGCAGGTCGCCGCGGTGGCCGTGGGCGTGGCCGTGCTCGTCCTGCTGCTGCTGCTCGGGCTGGGGGGCGCGCACTGCTACAGGACTAAGAAGTTGTCATCAAAAAATTTAAAGAATCCTTATGAAGAGCCAAGCAGAGAGGGTAGCAGTAGCAGGCCTTCAGACAGCGAGGCTAGGATGGCCTCTTTTCCCCAACCTTGGTTTGTGGTTATAAAGGAACATCAAAATCTCAGGAATGGAAGTCAACCTATGGCCCTCAAGGATGGTGAGTCAGCAGATGTTAGCCAATTTTCCTCTCCAGAGCCAGGGTCAGTAAAACGGACCTCATGGAGAAATGAACACCAGTTATATAAGGACACAGAGCAAGGCTGCTGCACTCCACCATCCAGTAATAGAGGCACCGGCTCTCAGTCAATGGAGCAGAGCTTTTCTGTCCCCTCCTATGAGGCACAGCCCATTGCTTTGGGGGTTGAGAAGCCACAGTCTCTCCTATCAGCTAACCCTTATTGCAACAAAGGGCCCCAGATCCACCACACCAAATGA

### Canine HBEGF

ATGAAGCTGCTGCGGTCAGTGGTGCTGAAGCTCTTTCTGGCTGCAGTGCTCTCGGCGTCGGTGACTGGCGAGAGCTTGGGGCGTCTTCGGAGAGGGCTGGCGGCCGGAACTGGCAACCCGGACTCTCCCACCGAATCCACGGACCGGCTGCTGCCCCCGGAAGGCGGCCGGGCCAGGGAAGTCCTGGACTTAGAAGAGACGGACCTGGACCTTTTAAGAGCAGCTGCTTTCTCCTCCAAGCCACAGGCTCTGGCCACACCCAGTAAGGAGGAACGTGGGAAAAAAAAGAAGAAAGGCAAAGGCTTAGGGAGGAAGAGAGACCCGTGTCTTCGGAAATACAAGGACTTCTGCATCCATGGAGAATGCAAATATGTGAAGGAGCTCCGGGCTCCATCCTGCATCTGCCACCCCGGTTACCATGGAGAGAGGTGCCATGGGCTGAGCCTTCCAGTAGAAAATCGCTTATATACTTACGACCATACAACCATCTTGGCTGTGGTGGCTGTGGTGCTGTCATCTGTCTGTCTGCTTGTCATCGTGGGGCTTCTCATGTTTAGGTACCATAGGAGAGGAGGTTATGATGTGGAAAGTGAAGAGAAAGTGAAGTTAGGCATGACTACTTCCCACTGA

### Canine EREG

ATGGAGCCGCGCCGCCTGCTGCTGTGCCTGGGTTTCCATCTTCTCCACGCGGTTCTCAGCACCACTGTGATTCCTTCCTGCATGCCGGGAGAATCCGAAGATAATTGCACGGCATTAGTTCAGATAGAAGACAACCCACGTGTGGCTCAAGTGTCAATAATAAAGTGTGGCTCTGACATGAATGGCTACTGTTTGCATGGACAATGCATCTACCTGGTGGACATGAGTCAAACGTACTGCAGGTGTGAAGTGGGTTACACTGGTGTCCGATGCGAGCACTTCTATTTAACTGTCCAACAGCCCTTGAGCAAAGAATATGTGGCTTTGACTGTGATTCTCATTATCTTGTTTCTTATCATAGTCGCCGGTTCCCTATACTACTTCTGCAGATGGTACAGAAATCGAAAAAGTAAAGAACCAAAGCAGGAATACAAAAGGGTGACGTCAGGGGATCCAGCATTGCCACAAGTCTGA

### Canine TGFα

ATGGTGCCCTCGGCCGGACGGCTCGCCCTGCTCGCGCTGGGTGTCCTGCTGGCCGCGGGCCAGGCCCTGGAGAACAGCACGTCCGCCCTGAGTGCCAGACCACCAGTGGCCGCTGCAGTAGTGTCTCATTTTAACGATTGCCCAGATTCCCATAGCCAGTTCTGCTTCCACGGGACTTGCAGATTTCTGGTTCAAGAAGATAAACCCGCTTGTGTATGCCACAGTGGCTATGTCGGGGCACGCTGTGAACACGCCGACCTTCTTGCTGTCGTAGCAGCGTCCCAAAAGAAGCAGGCCATCACTGCTCTGGTGGTGGTCTCTATCGTGGCGTTGGCCGTGCTGATTATCGCATGTGTTCTGATCCATTGCTGTCAGGTGCGCAAGCACTGTGAGTGGTGTCAGGCCCTCTTGTGTCGGCATGAGAAGCCCTCCGCATTGCTTAAGGGACGAGCTGCTTGCTGTCATAGCGAGACAGCCGTCTGA

### Canine NRG1

ATGGAACCCGACGCCAACAGCAGCAGTAGAGCCCCTGCTGCCTTTCGGGCTAGCTTCCCACCTCTGGAAACCGGCCGGAACCTGAAGAAAGAGGTGTCCAGAGTCCTGTGCAAGAGATGCGCCCTGCCTCCTCGGCTGAAAGAGATGAGAAGCCAAGAGTCTGCCGCCGGAAGCAAGCTGGTGCTGAGATGTGAAACCAGCAGCGAGTACAGCAGCCTGAAGTTCAAGTGGTTCAAGAACGGCAACGAGCTGAACCGGAAGAACAAGCCCCAGAACATCAAGATCCAGAAGAAGCCCGGCAAGAGCGAGCTGAGAATCAGCAAAGCCAGCCTGGCCGATAGCGGCGAGTACATGTGTAAAGTGACCTCCAAGCTGGGCAACGACAGCGCCAGCGCCAATATCACCATCGTGGACAGCAACGACATCATCACCGGCATGCCTGCCAGCACCGAGAGGGCTTATGTGTCTAGCGAGAGCCCCATCCGGATCAGCGTTAGCACAGAAGGCGCCAACACAAGCAGCAGCACCTCTACCAGCACCACCGGCACATCTCACCTGGTCAAGTGCGCCGAGAAAGAAAAGACCTTCTGCGTGAACGGCGGCGAGTGCTTCATGGTCAAGGACCTGAGCAACCCCAGCCGGTATCTGTGCAAGTGTCAGCCCGGTTTTACCGGCGCCAGATGCACCGAGAATGTGCCCATGAAGGTGCAGAATCAAGAGAAGGCCGAGGAACTGTACCAGAAACGGGTGCTGACCATCACAGGCATCTGTATCGCCCTGCTGGTCGTGGGCATTATGTGCGTGGTGGCCTACTGCAAGACCAAGAAGCAGCGGAAGAAGCTGCACGACCGGCTGAGACAGAGCCTGAGAAGCGAGAGAAACAACATGGTCAATATCGCCAACGGACCCCACCATCCTAATCCTCCACCTGAGAACGTGCAGCTGGTCAACCAGTACGTGTCCAAGAACGTGATCAGCTCCGAGCACATCGTGGAACGCGAGGCCGAGACAAGCTTTAGCACCAGCCACTACACCAGCACAGCCCACCACAGCACCACAGTGACACAGACCCCAAGCCACAGCTGGTCCAATGGCCACACCGAGTCCATCATCAGCGAGTCCCACAGCGTGATCATGATGAGCAGCGTGGAAAACAGCCGGCACTCTAGCCCTAGCGGAGGACCTAGAGGCAGACTGAATGGCCTCGGCGGACCCAGAGAGTGCAACAGCTTTCTGAGACACGCCAGAGAGACACCCGACAGCTACAGAGATAGCCCTCACAGCGAGAGATACGTGTCCGCCATGACAACCCCTGCCAGAATGAGCCCCGTGGACTTTCACACACCTAGCAGCCCTAAGAGCCCTCCTAGCGAGACATCCCCTCCAGTGTCTAGCACAACCGTGTCCATGCCTAGCATGGCCGTGTCACCCTTCGTGGAAGAGGAAAGACCTCTGCTGCTGGTCACCCCTCCTAGACTGAGAGAGAAGTACGATCACCACAGCCAGCAGTTCAACAGCTACCACCACAATCCTGCTCACGAGAGCAACAGCCTGCCTCCATCTCCTCTGAGAATCGTCGAGGACGAGGAATACGAGACAACCCAAGAGTACGAGCCCGCTCAAGAGCCCGTGAAGAAACTGACCAGCTCCAGACGGGCCAAGCGGACCAAGCCTAATGGACACATTGCCAACCGGCTGGAAATGGACTCCAATGCCTCTGCCGAGGGCACCAACAGCGAAAGCGAGACAGAGGATGAGAGAGTGGGCGAAGATACCCCATTCCTGGGCATCCAGAATCCTCTGGCCGCCTCTCTTGAAGCCGCTCCTGCTTTTAGACTGGCCGACAGCAGAACAAACCCAGCCGGCAGATTCAGCACCCAAGAGGAACTGCAGGCCAGACTGTCTAGCGTGATCGCCAACCAGGATCCTATCGCCGTCTGA
